# Endothelial cells secrete small extracellular vesicles bidirectionally containing distinct cargo to uniquely reprogram vascular cells in the circulation and vessel wall

**DOI:** 10.1101/2023.04.28.538787

**Authors:** Sneha Raju, Steven R. Botts, Mark Blaser, Kamalben Prajapati, Tse Wing Winnie Ho, Crizza Ching, Natalie J Galant, Lindsey Fiddes, Ruilin Wu, Cassandra L. Clift, Tan Pham, Warren L Lee, Sasha A Singh, Elena Aikawa, Jason E Fish, Kathryn L Howe

**Affiliations:** Toronto General Hospital Research Institute, University Health Network, Toronto, Canada.; Institute of Medical Science, University of Toronto, Toronto, Canada.; Division of Vascular Surgery, Toronto General Hospital, Toronto, Canada.; Faculty of Medicine, University of Toronto, Toronto ON, Canada.; Center for Interdisciplinary Cardiovascular Sciences, Cardiovascular Division, Department of Medicine, Brigham and Women’s Hospital, Harvard Medical School, Boston, MA, USA.; Keenan Research Centre for Biomedical Science, St. Michael’s Hospital, Toronto, ON, Canada.; Princess Margaret Cancer Center, Toronto, Canada.; Department of Laboratory Medicine and Pathobiology, University of Toronto, Toronto, Canada.; Center for Excellence in Vascular Biology, Cardiovascular Division, Department of Medicine, Brigham and Women’s Hospital, Harvard Medical School, Boston, MA, USA.; Peter Munk Cardiac Centre, Toronto General Hospital, Toronto, Canada Short Title: Directional endothelial cell-cell communication

**Keywords:** atherosclerosis, monocyte, vascular smooth muscle cell, RNAseq, proteomics

## Abstract

Rationale: Extracellular vesicles (EVs) contain bioactive cargo including microRNAs (miRNAs) and proteins that are released by cells as a form of cell-cell communication. Endothelial cells (ECs) form the innermost lining of all blood vessels and thereby interface with cells in the circulation as well as cells residing in the vascular wall. It is unknown whether ECs have the capacity to release EVs capable of governing recipient cells within two separate compartments, and how this is affected by endothelial activation commonly seen in atheroprone regions.

Objective: Given their boundary location, we propose that ECs utilize bidirectional release of distinct EV cargo in quiescent and activated states to communicate with cells within the circulation and blood vessel wall.

Methods and Results: EVs were isolated from primary human aortic endothelial cells (ECs) (+/- IL-1β activation), quantified, and analysed by miRNA transcriptomics and proteomics. Compared to quiescent ECs, activated ECs increased EV release, with miRNA and protein cargo that were related to atherosclerosis. RNA sequencing of EV-treated monocytes and smooth muscle cells (SMCs) revealed that EVs from activated ECs altered pathways that were pro-inflammatory and atherogenic. Apical and basolateral EV release was assessed using ECs on transwells. ECs released more EVs apically, which increased with activation. Apical and basolateral EV cargo contained distinct transcriptomes and proteomes that were altered by EC activation. Notably, basolateral EC-EVs displayed greater changes in the EV secretome, with pathways specific to atherosclerosis. *In silico* analysis determined that compartment-specific cargo released by the apical and basolateral surfaces of ECs can reprogram monocytes and SMCs, respectively.

Conclusions: The demonstration that ECs are capable of polarized EV cargo loading and directional EV secretion reveals a novel paradigm for endothelial communication, which may ultimately enhance our ability to design endothelial-based therapeutics for cardiovascular diseases such as atherosclerosis where ECs are persistently activated.

**Non-standard Abbreviations and Acronyms:**

cryo-EM: cryogenic electron microscopy
EC: endothelial cell
EV: extracellular vesicle
GO: gene ontology
HAEC: human aortic endothelial cell
SMC: human aortic vascular smooth muscle cell
IL-1β: interleukin 1 beta
KEGG: Kyoto encyclopedia of genes and genomes
LC-MS: label-free liquid-chromatography mass spectrometry
MVB: multivesicular body
miRNA: microRNA
RNAseq: RNA sequencing
TEM: transmission electron microscopy
TIRF: total interal reflection fluorescence microscopy
miRNA: microRNA

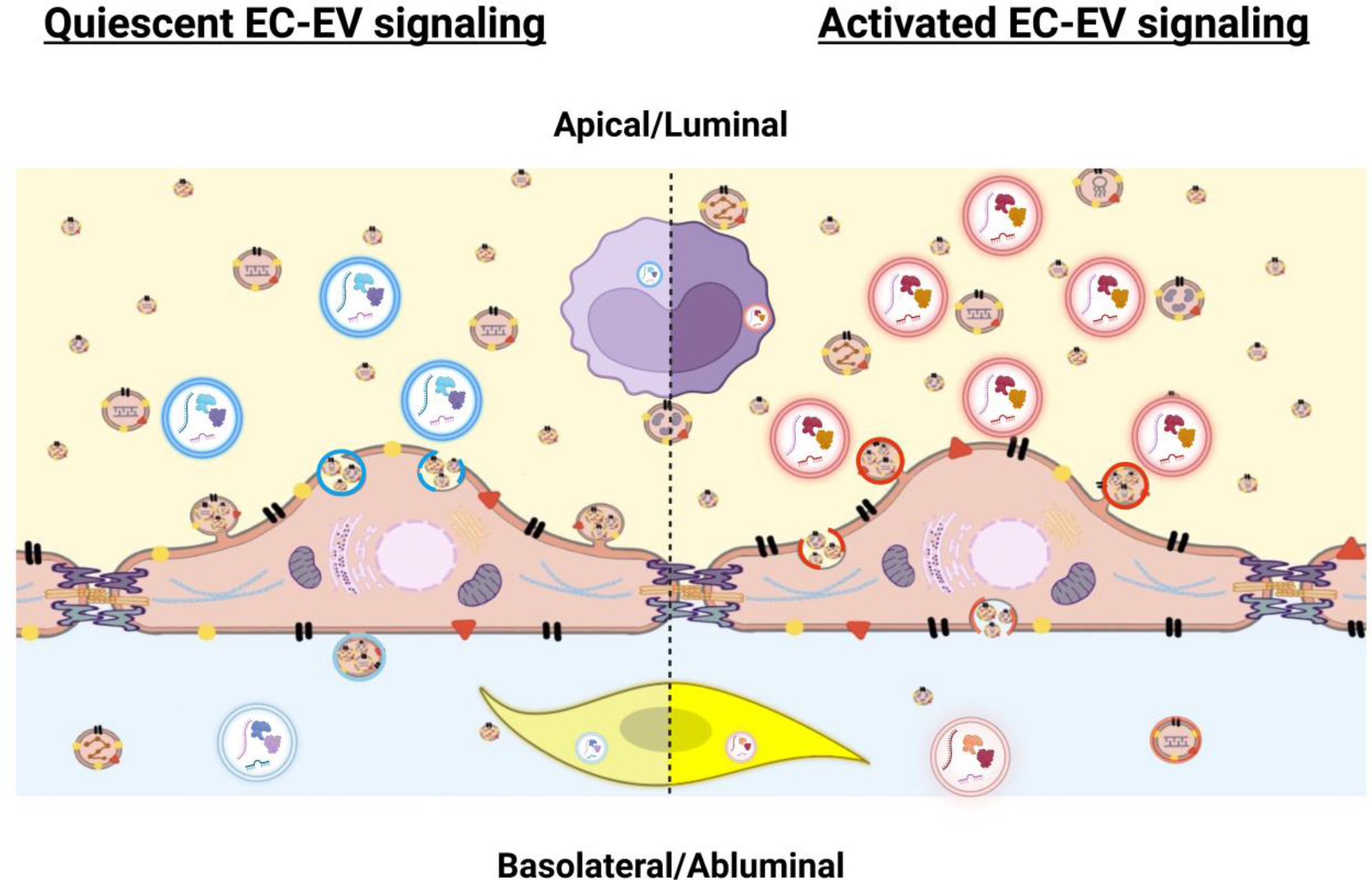

**Graphical abstract: Polarized endothelial extracellular vesicle communication with luminal and abluminal vascular cells:** Endothelial cell small extracellular vesicle (EC-EV) release from apical (luminal) and basolateral (abluminal) surfaces in quiescence and after endothelial activation. Quiescent EC-EVs are depicted in blue (bright blue=apical, light blue=basolateral), while activated EC-EVs are depicted in red (bright red=apical, light red=basolateral). Luminal monocyte is represented in purple with upregulation of pro-inflammatory transcripts (bright purple) after uptake of activated EC-EVs from the apical surface, compared to uptake of quiescent apical EC-EVs (light purple). Basolateral EC-EVs are taken up by an abluminal resident smooth muscle cell depicted in yellow. Smooth muscle cell uptake of activated basolateral EC-EVs with upregulation of pro-inflammatory/pro-atherogenic transcripts (bright yellow), as compared to uptake of quiescent EC-EVs (light yellow).

## Introduction

Endothelial cells (ECs) line the entire vasculature, forming the largest distributed organ in the human body with a unique secretome that maintains vascular health and regulates disease pathogenesis.^1, 2^ Given their critical interface between circulating blood and the vascular wall, ECs have the potential to communicate with both luminal and abluminal cells.^3^ Dysfunctional endothelium, as seen in chronic diseases such as atherosclerosis, is one of the earliest detectable pathophysiological events and contributes directly to the development of a wide range of cardiovascular diseases.^4–7^ Specifically, EC-based intercellular communication with circulating cells and resident vascular cells lies at the core of atherogenesis.^3^ Extracellular vesicles (EVs) have emerged as important mediators of cell-cell communication.^8, 9^ Cells produce EVs with specific cargo, such as microRNA (miRNA) and protein, based on their physiologic or pathologic state, and the contents of EVs can be delivered to recipient cells to mediate biologic effects.^9, 10^ Under quiescent states, cultured ECs release EVs that tame monocyte activation^11^ and provide atheroprotective communication to smooth muscle cells^11, 12^, while EVs derived from activated endothelium stimulate monocyte adhesion^13, 14^ and promote monocyte differentiation into inflammatory macrophages^15^. Although EC-EV based cell-cell communication is emerging as an important vector mediating multicellular disease states, it remains unresolved whether ECs can direct EV release in a polarized fashion to communicate separately with luminal and abluminal cells.

ECs are uniquely situated at the interface of the blood vessel lumen and wall where they are positioned in an asymmetrical extracellular environment. In endothelial biology there is a precedent for polarized structural and functional arrangements. EC proteins are polarized to luminal (apical) and abluminal (basolateral) surfaces to facilitate distinct functions (e.g., glycocalyx versus cell adhesion).^16^ In cardiovascular disease, endothelial polarity proteins (e.g., Scrib) play key roles in establishing endothelial identity and are atheroprotective.^17^ Given their location and evidence of apical-basal polarity, it is conceivable that ECs secrete EVs in a polarized manner and alter cargo based on environmental cues as a mechanism for distinct communication.^18^ Delineating polarized endothelial EV communication would fundamentally alter the approach to vascular biology and cardiovascular disease.

In the current study, we demonstrate the crucial role for EC-based communication in cardiovascular disease conditions using primary human aortic endothelial cells (HAECs) exposed to IL-1β, a key cytokine increased in relation to disease severity in patients with atherosclerosis^19^ and the focus of the CANTOS trial that definitively proved the role of inflammation in cardiovascular events.^20, 21^ We determined that HAECs increase EV release when activated with IL-1β and package miRNA and protein cargo that is clearly distinct from the quiescent state. Notably, these EVs have functional effects, with cellular reprogramming of primary human monocytes and human aortic smooth muscle cells (SMCs) as determined by RNA-sequencing. Using multiple lines of evidence, we demonstrate that ECs are capable of directional (i.e., apical and basolateral) EV release in both quiescent and activated states, providing a mechanism for endothelial communication strategies with cells in separate extracellular compartments. Moreover, ECs shuttle distinct EV-cargo to the luminal and abluminal compartment that is altered upon activation. *In silico* analysis further determined that polarized, compartment-specific EC-EVs have the capacity to communicate with monocytes and SMCs - cells found in the luminal and abluminal compartments respectively - to instigate unique changes in key pro-atherogenic transcripts and pathways.

## Methods

Detailed Materials and Methods are available in the Data Supplement. The authors declare that all supporting data are available within the Data Supplement.

## Results

### A subpopulation of small EVs is increased after EC activation

‘EV’ is a broad term for bilayered nanoparticles that are secreted via several routes, including plasma membrane-derived microparticles and small EVs (sEVs; also known as exosomes), which are secreted from a specialized subset of endosomes called multivesicular bodies (MVBs).^22^ sEVs were enriched via serial ultracentrifugation (Online Figure I A) from media of unstimulated (quiescent) or activated primary human aortic endothelial cells (HAECs) (IL-ip, 100 pg/mL, 24 h; Online Figure I B-C). Activated HAECs released more sEVs (size range 30­200 nm) than quiescent HAECs, as measured by nanoparticle tracking analysis (NTA) (Figure 1A, B). To confirm that these nanoparticles were EVs, we performed western blot analysis that showed expression of common EV markers, including CD63, Alix, and CD9 (Figure 1C), but only CD63-positive sEVs were significantly increased by endothelial activation (Figure 1D and Online Figure I D). No morphological or size differences between sEV isolates from quiescent and activated HAECs were noted using cryo-electron microscopy (cryo-EM) or NTA (Figure 1E-F). However, TEM of activated HAECs *in situ* demonstrated increased MVBs positioned near the cell surface compared to quiescent cells (Figure 1G). Increased sEV release by activated ECs was not accompanied by increased EV biogenesis markers (e.g., TSG101, Caveolin, Flot1) at the mRNA or protein level - consistent with publicly available data (GEO accession: GSE89970 (Online Figure II)).^23^ Together, these results demonstrated the release of CD63­positive endothelial sEVs is significantly increased in response to a cardiovascular disease-related stimulus and motivated us to determine whether differential EV cargo might accompany this altered EV landscape.

**Figure 1.**
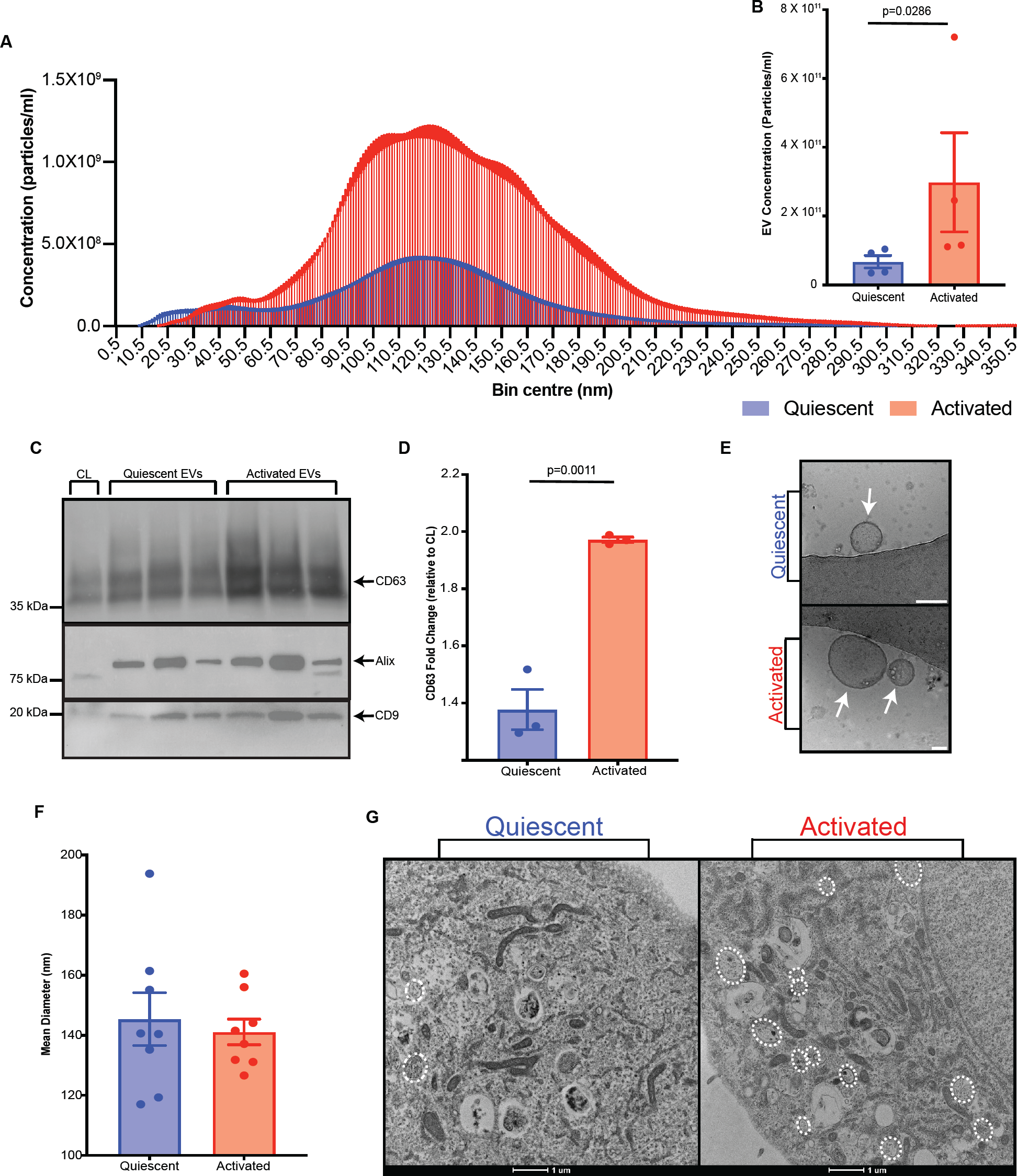
Endothelial cells release increased CD63-positive sEVs in response to activation. **A,** Nanoparticle tracking analysis (NTA) of EV concentration binned by particle size after isolation from HAEC conditioned media (8 X 10^7^ cells, from quiescent (EV-free media, 24 h) and activated (100 pg/mL IL-1β in EV-free media, 24 h) states (*n*=3-8). **B**, Quantification of EC-EV mean concentration across all EV sizes. **C**, Western blot depicting EV markers (CD63, Alix, and CD9) in EV lysates isolated from supernatants of quiescent and activated HAECs and HAEC cell lysate (CL) control. Arrows show position of correct protein band and molecular weights markers indicated on left. **D**, Densitometry of EV lysate derived CD63 normalized to HAEC cell lysate control. **E**, Cryo-EM of EVs isolated from quiescent and activated HAEC cell supernatant. Arrows indicate EV structures. Scale bar=50 nm. **F**, Quantification of EV mean diameter by NTA. **G**, Transmission electron microscopy of 90 nm ultramicrotomed HAEC monolayers. Dashed circles indicate multivesicular bodies. Representative image (n=3). Bar graphs show mean ± SEM. Statistical significance assessed by Mann-Whitney test (B) and unpaired *t* test (D, F).

### sEV cargo displays a pro-atherogenic signature after EC activation

EV cargo including miRNA and proteins mediate biological effects. Next generation miRNA sequencing of HAEC sEVs showed independent clustering of sEV-miRNA from activated versus quiescent states with greater heterogeneity of sEV-miRNA expression among quiescent ECs (Figure 2A). sEVs from quiescent and activated ECs have similar distributions of miRNA abundance (i.e., normalized miRNA counts), but EC activation drives sEV-miRNA cargo towards a more homogenous transcriptome (i.e., reduces variation in miRNA counts between samples) (Figure 2B). MiRNA cargo is differentially expressed in EC-sEVs, with 192 transcripts increased in activated conditions and 305 transcripts in quiescent EC-sEVs (Figure 2C). Several known endothelial-enriched miRNAs such as miRNA-126, miRNA-92a, and miRNA-181, were abundantly expressed in both conditions (Online Figure III A). To assess the functional implications of altered sEV-miRNA cargo, we employed KEGG pathway analysis of differentially expressed sEV-miRNAs in quiescent conditions (Online Figure III B). MiRNA-513a-3p, miRNA-208b-3p, and miRNA-587 were found to govern pathways involved in cell-cell communication, cell cycle and metabolism, and cell signaling (Figure 2D, Online Figure III D). Conversely, several regulators of inflammation such as miRNA-146a-5p, miRNA-146b-3p, and miRNA-98-5p were increased in activated endothelial sEVs (Online Figure III C), governing proatherogenic pathways involved in inflammatory signaling, cell death and clearance, cell matrix interactions and cell-cell communication with circulating and resident vascular and inflammatory cells (Figure 2E, Online Figure III E).

**Figure 2.**
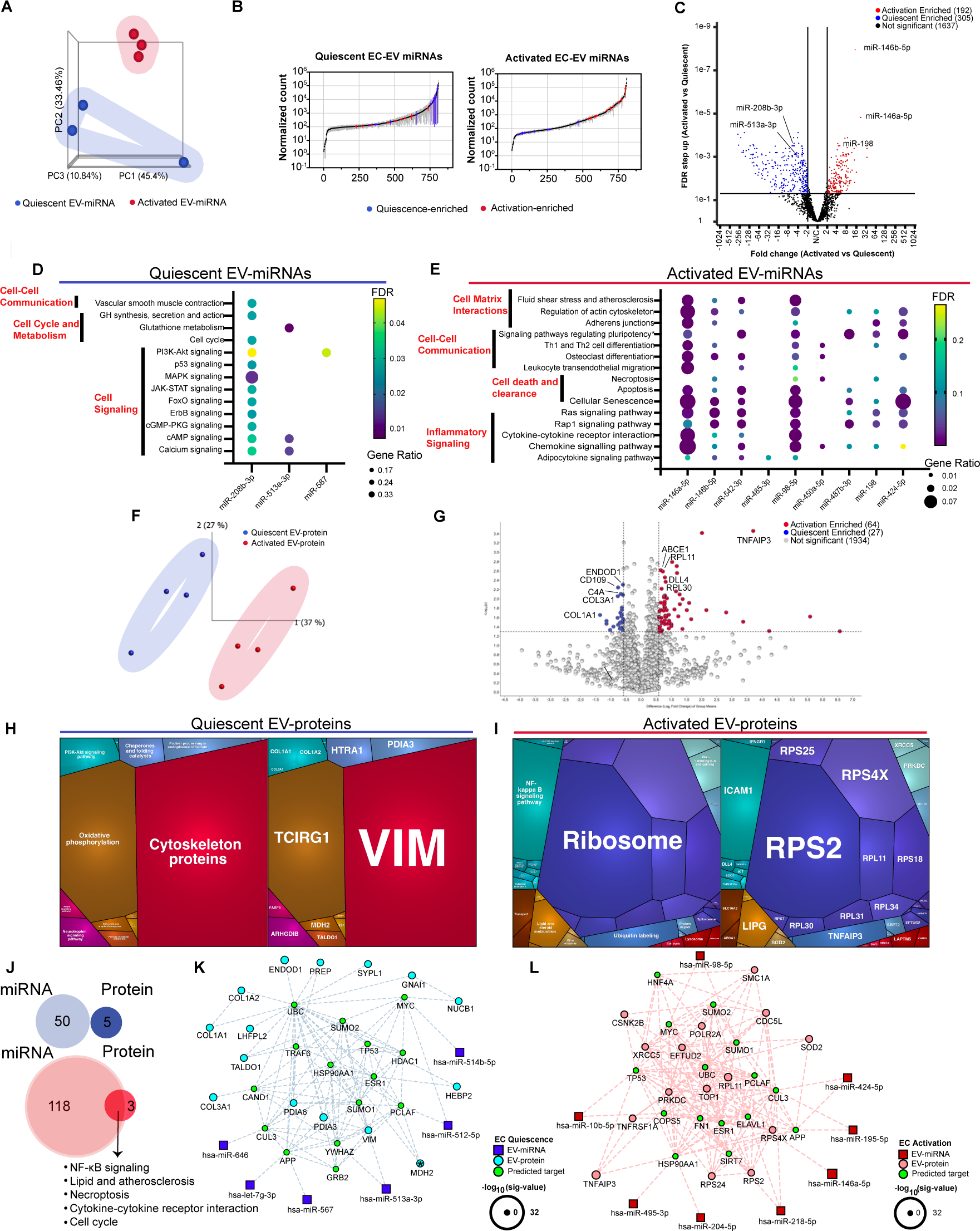
Endothelial sEV miRNA and protein cargo are distinct in identity and predicted function in activated versus quiescent conditions. **A,** Unfiltered principal component analysis (PCA) showing miRNA profiles of sEVs isolated from conditioned media of activated (red) versus quiescent (blue) HAECs (8 X 10^7^ cells, 100 pg/mL IL-1β, 24 h). **B,** Rank plots using normalized counts (arithmetic mean + SEM). Top 10 activation-and quiescence-enriched miRNAs are highlighted in red and blue, respectively. **C,** Volcano plot of HAEC secreted EV miRNA transcriptome with red and blue representing EV-miRNA contents enriched in activated and quiescent states, respectively (FDR step up < 0.05, Fold Change |2|) **D,** Pathway analysis of top 10 (by FDR) quiescent HAEC-EV enriched miRNAs (miRTarBase) delineated significant KEGG pathways (FDR < 0.05) for miRNA associations of miR-208b-3p, miR-513a-3p, and miR-587. Data points are sized by GeneRatio (genes altered in pathway/total number of unique genes in analysis) and colour-scaled by FDR. **E,** Pathway analysis of top 10 (by FDR) activated HAEC-EV enriched miRNAs (miRTarBase). Shown are individual miRNA associations of KEGG pathways of interest. Data points are sized by GeneRatio (genes altered in pathway/total number of unique genes in analysis) and colour-scaled by FDR. **F,** Unfiltered PCA showing protein profiles of sEVs isolated from conditioned media of activated (red) versus quiescent (blue) HAECs as in (**A). G,** Volcano plot of HAEC secreted EV proteome with red and blue representing EV-protein contents enriched in activated and quiescent states, respectively (p< 0.05, Fold Change |1.5|). **H,** Proteomap (v2.0, Homo Sapiens) generated from all differentially enriched quiescent EC-EV proteins weighted by mass abundance. KEGG orthology terms (left) and respective proteins (right) contributing to the pathways are illustrated. **I,** Proteomap (v2.0, Homo Sapiens) generated from all differentially enriched activated EC-EV proteins calculated as in **(H). J,** Overlapping KEGG pathways between the top 10 (by FDR) differentially enriched EV-proteins and all differentially enriched EV-proteins in quiescent (top, blue) and activated states (bottom, red). **K-L,** EV interactome generated by capturing differentially expressed EV-miRNA (top 25 by FDR) and all EV-proteins in quiescent **(K)** and activated **(L)** states, followed by network reduction to retain the top 15 of each group based on degree of interactions. EV-miRNA shown in blue or red, EV-proteins in turquoise or pink, and predicted targets in green. Node size denotes significant value. Data shown represent n=3-4 independent experiments. Cancer-, and infection-associated pathways were excluded from analysis. Data is represented as mean ± SEM.

Protein cargo of endothelial sEVs was assessed via label-free liquid-chromatography mass spectrometry (LC-MS) and yielded several known EV markers (Online Figure IV A-B). Similar to sEV-transcriptomics, distinct protein profiles were identified in activated versus quiescent HAEC sEVs (Figure 2F). There were 27 and 64 EC-sEV proteins that were differentially enriched in quiescent and activated states, respectively (Figure 2G). Quiescent EC-sEVs contained several abundantly enriched proteins with homeostatic roles in SMCs and extracellular matrix production (e.g., COL3A1, COL1A2, COL1A1), vascular endothelial maintenance (e.g., ENG, VIM) and cell metabolism (TCIRG1, MDH2, TALDO1, MIF, MBOAT7) (Figure 2H). Activated EC-sEVs contained key drivers of atherosclerosis including adhesion molecules (e.g., ICAM-1), inflammatory and stress proteins (e.g., IL1RAP, IFNGR1, TNFRSF1A, DLL4, SOD2), ribosomal proteins (e.g., RPS25, RPL11, RPL30, RPL34), and lipoprotein metabolism (LIPG), (Figure 2I and Online Figure IV C). KEGG pathway analysis of the sEV proteome in activated HAECs delineated five key pathways that were shared with predicted targets of EC-sEV miRNAs -NF- kB signaling, lipid and atherosclerosis, necroptosis, cytokine-cytokine receptor interaction, and cell cycle - demonstrating the shift in the sEV secretome towards an atherogenic payload (Figure 2J). Given sEV-derived miRNAs and proteins may function collectively, we assessed their interactions with predicted gene targets and generated sEV interactomes (Figure 2 K-L). Predicted targets of sEV cargo from the quiescent endothelial state included genes involved in protein folding and degradation, DNA regulation and repair, and cell proliferation and differentiation (Figure 2K). The activated sEV network appeared denser than its quiescent counterpart and while there was overlap between predicted targets of activated and quiescent EC-sEVs, activated EC-sEVs altered genes involved in transcriptional and translational regulation (Figure 2L). These data show that sEV cargo and release is substantially altered upon endothelial activation and has the potential to impact communication with surrounding cells to potentiate inflammatory messaging.

### Activated EC-sEVs uniquely reprogram circulating monocytes and resident SMCs

Given activated ECs released distinct sEV cargo that reflected pro-inflammatory and atherosclerosis-relevant pathways, we next determined the direct effects of these EC-sEVs on human monocytes or SMCs - key cells involved in the development of chronic vascular diseases such as atherosclerosis. Human primary CD14+ monocytes were treated for 24 hours with sEVs isolated from activated or quiescent HAECs (Online Figure V A). Monocyte RNA was isolated, and RNA sequencing (RNAseq) was performed to delineate the cellular response of monocytes to EC-sEVs. Principal component analysis of RNAseq revealed distinct profiles for monocytes exposed to quiescent EC-sEVs, activated EC-sEVs, and cell culture media-alone control (Figure 3A). Heatmap analysis showed groups cluster separately and that most transcripts lay within protein coding regions (Online Figure V B). While sEVs from activated ECs had biological effects on monocytes when compared to media-alone control (Online Figure V C), more remarkable was the observation that sEVs from activated ECs drove unique biological activity on monocytes when compared to monocytes exposed to sEVs from quiescent ECs (Figure 3B). To better understand the cellular reprogramming being initiated in monocytes, pathway analysis was performed. As seen in Online Figure VD, monocytes exposed to activated EC-sEVs versus media-alone control significantly upregulated pathways in migration and chemotaxis, inflammation, and vascular development. To compare the specific impact of endothelial activation and altered sEV cargo on recipient monocytes, we analyzed monocyte responses to activated EC-sEVs versus quiescent EC-sEVs (Figure 3C): notably, pathways involved in adhesion and migration, inflammation, proliferation and differentiation, and apoptosis emerged. Given our well-characterized data from EC-sEV transcriptomics and proteomics, we integrated sEV cargo data with our monocyte recipient cell RNAseq to create an interactome that delineated differentially expressed transcripts that were likely regulated by EC-sEVs (Figure 3D-E). Five transcripts - down-regulation of monocyte SOX4, HIF1A, SOD2, TNFAIP3 (Figure 3E) and upregulation of monocyte LMNB1 (Figure 3D) by activated EC-sEVs - regulated the pathways highlighted in red (Figure 3C), including inflammatory signalling with IL-12 and TNF, and regulation of leukocyte proliferation and apoptosis. Overall, there was a strong intersection of the sEV-miRNAome, sEV-proteome, and recipient monocyte transcriptome centering on oxidative stress (e.g., HIF1A, SOD2, GNL3) under activated EC conditions (Figure 3D-E). Together, these data demonstrate that several pathways identified from miRNA and proteomic analysis of EC-sEVs are shared with pathways found in EV-treated monocytes, and that sEVs from an activated endothelium have specific biologic effects on recipient cells that differ from those derived from the quiescent state.

**Figure 3.**
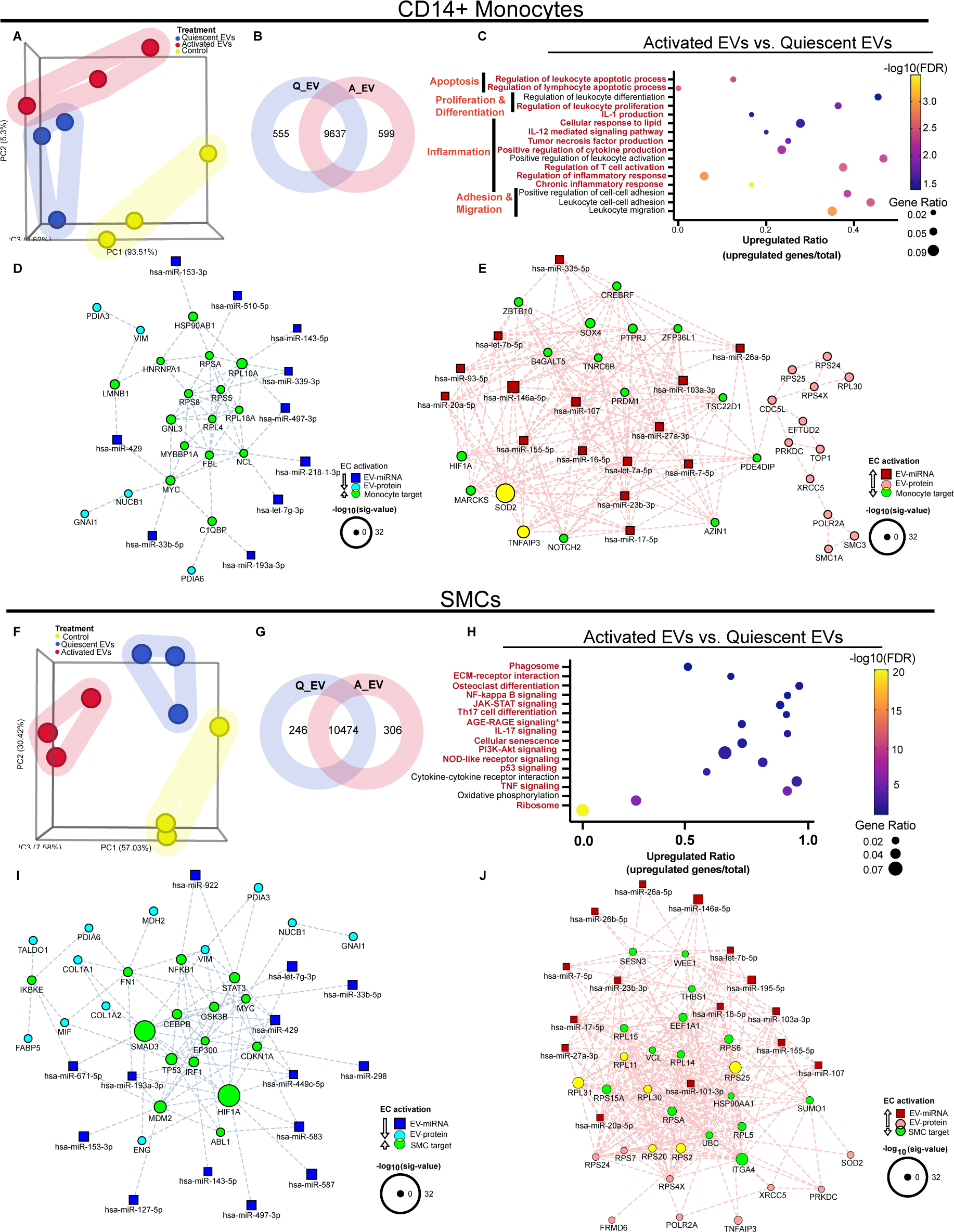
Endothelial sEVs distinctly alter the transcriptional landscape of recipient monocytes and smooth muscle cells depending on whether they are derived from quiescent or activated endothelium. **A,** Unfiltered PCA plot depicting clustering of media control (yellow), quiescent EC-EVs (blue), and activated EC-EVs (red) treated CD14+ monocyte mRNA transcriptome (*n*=3). **B,** VENN diagram depicting number of shared and unique RNA transcripts in comparison of activated vs quiescent EC-EV treatment. **C,** GO pathway analysis of the effects of activated versus quiescent EC-EVs on the monocyte RNA transcriptome (adjusted p-values <0.05 and |log2(FoldChange)| >0). Data points are sized by GeneRatio (genes altered in pathway/total number of unique genes in analysis) and colour-scaled by FDR. Upregulated ratio was calculated by dividing the number of upregulated genes by the total number of genes known to function in each pathway. **D-E,** Interactomes integrating activated EC-EV secretome (top 15 miRNAs and all EV-proteins) with differentially expressed monocyte transcripts based on degree of interactions. Downregulated EC-EV cargo and concordant upregulated monocyte targets are depicted in **(D)**. Upregulated EC-EV cargo and concordant downregulated monocyte targets are depicted in **(E). F,** Unfiltered PCA plot depicting clustering of EC-EV treated SMC mRNA transcriptome as in **A** (*n*=3). **G,** VENN diagram depicting SMC RNA transcripts as in **(B). H,** KEGG pathway analysis of the effects of activated versus quiescent EC-EVs on the SMC RNA transcriptome (adjusted p-values <0.05 and |log2(FoldChange)| >0). Data visualization completed as in **(C). I-J,** Interactomes as in **(D-E)** integrating differentially expressed SMC transcripts.

Since HAECs can communicate with luminal cells such as primary human monocytes via sEVs, we next assessed whether EC-sEVs can also regulate abluminal cells such as SMCs (Online Figure V A). Principal component and heat map analysis demonstrated that primary human aortic SMCs exposed to media-alone control, quiescent EC-sEVs, and activated EC-sEVs cluster distinctly with differentially expressed transcripts predominantly found within protein coding regions (Figure 3F, Online Figure V E). Similar to our observations with monocytes, sEVs from activated ECs had biological effect on SMCs when compared to media-alone control with 324 and 303 uniquely regulated transcripts, respectively (Online Figure V F). Exposure of SMCs to activated versus quiescent EC-sEVs led to shared, as well as distinct, transcript changes (Figure 3G). While there were clearly shared pathways including ribosome, cellular senescence, and several inflammatory signaling pathways, they varied by importance as noted by FDR value, with reduction of ribosomal pathways dominating the activated EC-sEV treated SMCs (Figure 3H, Online Figure V G). Unique SMC pathways regulated by activated versus quiescent EC-sEVs included phagosome and p53 signaling (Figure 3H). Interactomes created by integrating sEV-secretome and recipient SMC transcriptomic data identified several predicted targets of sEV-miRNA and proteins that are involved in pathways highlighted in red (Figure 3H). Notably, activated EC-sEVs led to a decrease in several ribosomal (RPL15, RPSA, RPL5) and protein metabolism related transcripts (EEF1A1, UBC, HSP90AA1, SUMO1), while mediators of inflammatory signaling (NFKB1, IKBKE, HIF1A, IRF1, SMAD3, FN1) and cell cycle regulators (TP53, CEBPB, MYC, MDM2) were increased (Figure 3I-J). These data delineate several key pathways and transcripts modulated by EC-sEVs demonstrating their ability to communicate with SMCs and drive pro-inflammatory/pro-atherogenic changes in recipient cells. Taken together, these findings underscore the significant biological potential of sEVs released by an activated endothelium, with increased concentration and altered cargo capable of differentially affecting cells in the circulation (e.g., circulating monocytes, apical/luminal) or cells that reside in the vessel wall (e.g., SMCs, basolateral/abluminal). The directional potential for EC-sEVs as a mechanism for cellular communication to discrete compartments has not previously been explored.

### ECs can direct cell-cell communication via polarized release of sEVs

Given that quiescent and activated EC-derived sEVs have distinct transcriptomic and proteomic signatures with the capacity to alter monocyte and SMC function, we sought to determine whether ECs release sEVs in a polarized fashion where they might participate in divergent cell­cell communication strategies from their apical (luminal) and basolateral (abluminal) surfaces. To explore this concept, we employed a transwell system.^18, 24^ Maintenance of physiologic barrier function was confirmed by VE-cadherin localization to adherens junctions and 30 nm gold nanoparticle challenge (representing the smallest sEV) across EC monolayers (Online Figure VI A-B). Using transwells to enrich EVs from the media in upper (apical/luminal) and lower (basolateral/abluminal) chambers pushed the limits of low input analyses. We therefore employed ultracentrifugation for our initial sEV observations (NTA, cryo-EM, and EV-marker analysis by western blot) and then progressed to size exclusion chromatography for activation studies, and transcriptomic and proteomic analysis (Figure 4A, Online Figure IV D-E). sEVs were identified in isolates from apical and basolateral compartments and visualized as bilayered nanoparticles with dense cores (Figure 4B). Apical sEVs were larger than basolateral sEVs (Figure 4C). HAECs released more sEVs to the apical compartment as determined via nanoparticle counts (NTA) and western blot analysis of common EV markers, with a significant increase detected in CD63-positive sEVs (Figure 4D-F). To assess whether this property of polarized EC-sEV release was broadly applicable to other EC types, we recapitulated the findings with pooled human umbilical vein ECs (HUVECs) (Online Figure VI C-G). Given the novelty of basolateral EC-sEV release, we employed total internal reflection fluorescence (TIRF) microscopy to visualize sEV release from the basolateral surface of HAECs transiently transfected with pHluorin-CD63 plasmid.^25^ EVs were visualized when MVBs fuse with the plasma membrane and release sEVs into the neutral extracellular milieu. Their release was increased upon stimulation with histamine (Figure 4G-H) and was significantly inhibited upon addition of an EV release inhibitor, GW4869 (Figure 4I-J). We further delineated the role of polarized EV release in cell-cell communication by performing a quantitative assessment of EC-EV transfer to monocytes (Figure 4K). Using ECs transfected with exogenous *C. elegans* miRNA-39, ECs transferred miRNA-39 to monocytes in apical and basolateral compartments, with apical monocytes receiving most of the miRNA (Figure 4K). At the ultrastructural level, TEM images suggested that sEVs were contained within MVBs and were poised for release at both the apical and basolateral surfaces (Figure 4L). Given our data identified polarized sEV release by the endothelium as a putative mechanism for communication with circulating and resident vascular cells, we questioned whether ECs release compartment-specific cargo to communicate differentially with vascular cells.

**Figure 4.**
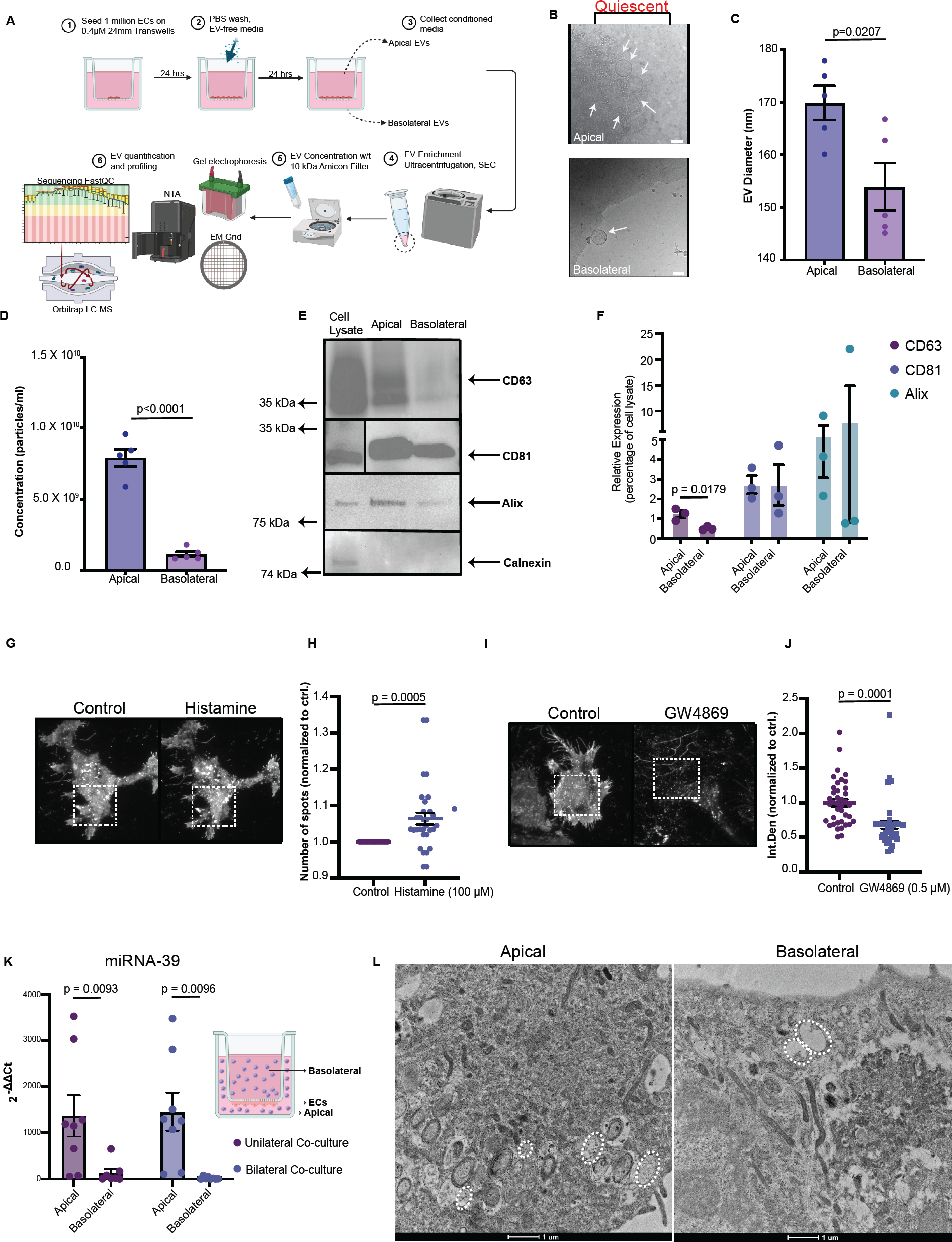
Multi-modal evidence determining quiescent endothelial cells release sEVs to apical and basolateral compartments. **A,** Workflow showing EC-EV isolation from compartments. Briefly, HAECs were seeded at confluence on semi-permeable transwell inserts to sequester EVs from apical and basolateral compartments. EVs were isolated by ultracentrifugation or size exclusion chromatography, concentrated, and validated according to MISEV2018 guidelines. Created with BioRender.com. **B,** Cryo-EM of representative images of apical (top panel) and basolateral (bottom panel) quiescent EC-EVs. Arrows denote EV structures. Scale bar 50 nm. **C-D,** Nanoparticle tracking analysis quantifying the mean diameter **(C)** and concentration **(D)** of EC-sEVs in apical and basolateral compartments. **E-F,** Western blot depicting protein expression of EV markers ((positive (CD63, CD81, Alix) and negative (Calnexin)), in cell lysate, apical EV and basolateral EV samples (**E)**. Arrows show position of correct protein band and molecular weights markers indicated on left. Densitometric analysis of EV markers **(F). G-J,** Total internal reflection fluorescence (TIRF) microscopy. Panels depicting ECs transfected with fluorescent plasmid (pHluorinCD63) set for detection of basolateral EV release +/- positive (histamine, 100 mM, 1 min) and negative (GW4869, 0.5 mM, 4 h) controls **(G, I)**. Quantification of basolateral EV release **(H, J)**. For histamine stimulated cells, vesicles in the TIRF zone were quantified and normalized to the number of cells in the field **(H)**. For GW4869 stimulated cells, integrated densities of CD63-pHluorin under basal conditions and after pre-treatment was quantified **(J). K,** Model for exogenous miRNA transfer between ECs and monocytes (see methods for full details). Briefly, HAECs were transfected with exogenous miRNA-39 (*C. elegans*) and then seeded onto an inverted transwell to avoid direct cell-cell contact with non-adherent monocytes. Monocytes were then placed either in a solitary chamber (apical or basolateral, unilateral co-culture experiment) or simultaneously in the apical and basolateral chambers (bilateral co-culture experiment), with monocytes harvested after 24 h, and RNA isolated to quantify miRNA-39 expression by RT-qPCR. **L,** Transmission electron microscopy of 90 nm ultramicrotomed HAEC monolayers. Embedded blocks were cut from the basolateral surface: the first 5 mm of resin cut was discarded to get to the apical surface. Circles indicate multivesicular bodies. Data shown represent n=3-4 independent experiments. Bar graphs show mean ± SEM. Statistical significance assessed by unpaired *t* test **(C,D,F,J)**, paired *t* test **(H)** and Mann-Whitney test **(K)** when data was not normally distributed.

### ECs release sEVs with distinct cargo to apical and basolateral compartments

Endothelial sEVs from the apical and basolateral compartments of quiescent and activated cells underwent next-generation miRNA sequencing and proteomic analysis via LC-MS (Figures 5-6). sEV miRNA cargo from quiescent ECs readily clustered by polarity (broad circles) with distinct transcriptomes observed between apical versus basolateral sEV collection (Figure 5A). This polarization of sEV cargo was preserved with EC activation (Figure 5A, Online Figure VII A). Under quiescent conditions, endothelial-specific miRNA were polarized in their secretion (e.g., miRNA-126 ^26, 27^ family apically and miRNA-144^28, 29^ family basolaterally; Figure 5B, left panel). This polarization was maintained upon EC activation, suggesting that these miRNAs may be constitutively released in a polarized fashion by ECs even under inflammatory stimuli (Online Figure VII B, right panel). Pathway analysis of sEV-miRNA from apical and basolateral compartments highlighted distinct compartment-specific pathways. In quiescent states, there were 319 and 145 distinct pathways predicted to be regulated by apical and basolateral sEV-miRNA, respectively. Among the top pathways, those related to apoptosis, p53 signaling, and chemokine signaling were enriched apically, while pathways related to cellular senescence, and lipid and atherosclerosis were modulated basolaterally (Figure 5B, right panel). Notably, the polarization of the EV-miRNA transcriptome and downstream pathways was maintained in activated states (Online Figure VII B, left panel), suggesting the importance of distinct directional communication in health and disease.

**Figure 5.**
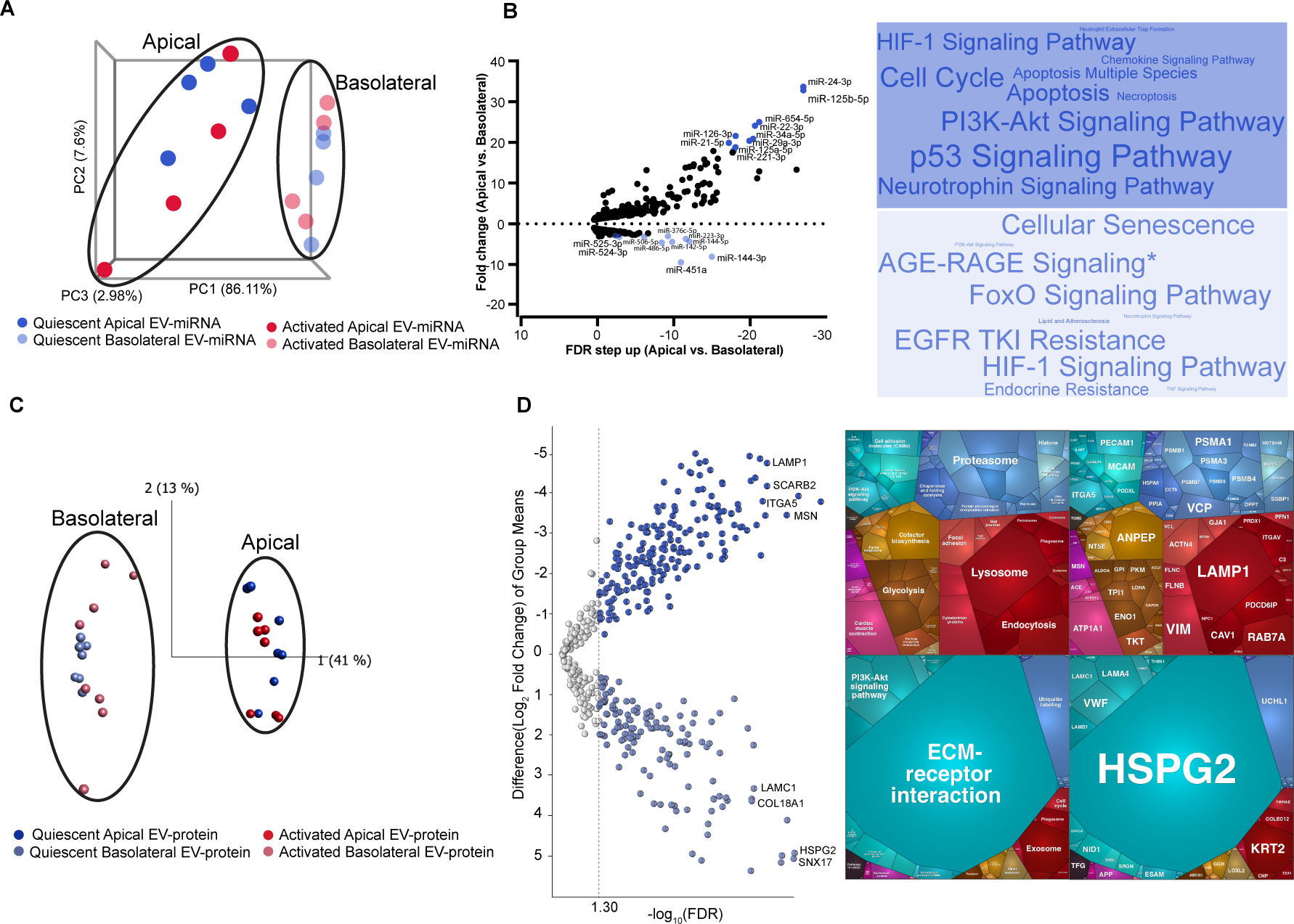
Quiescent endothelial cells release sEVs containing distinct miRNA and protein cargo to apical and basolateral compartments. **A,** Unfiltered PCA analysis of apical (dark colours) and basolateral (light colours) EV-miRNA depict clustering by polarity (broad circles). **B,** Volcano plot (left panel) of quiescent HAEC secreted EV-miRNA transcriptome enriched in apical (dark shading) versus basolateral (light shading) compartments. Top miRNA, by FDR step up, are labelled in each condition and used for downstream pathway analysis (FDR step up < 0.05). KEGG pathway analysis of labelled miRNA in each condition (FDR < 0.05), weighted by number of miRNAs participating in each pathway depicted by Word Cloud (right panel). **C,** Unfiltered PCA analysis of apical (dark colours) and basolateral (light colours) EV-protein profiles showing clustering by polarity (broad circles). **D,** Volcano plot (left panel) of quiescent HAEC secreted EV-proteome enriched in apical (dark shading) versus basolateral (light shading) compartments (FDR<0.05). All differentially enriched (FDR<0.05) apical versus basolateral proteins in quiescent conditions were inputted to generate proteomaps (v2.0, Homo Sapiens), weighted by protein mass abundance. Apical and basolateral proteomaps are represented by top and bottom panels, respectively. KEGG orthology terms (left) and respective proteins (right) contributing to the pathways are illustrated. *AGE-RAGE signaling in diabetic complications.

**Figure 6.**
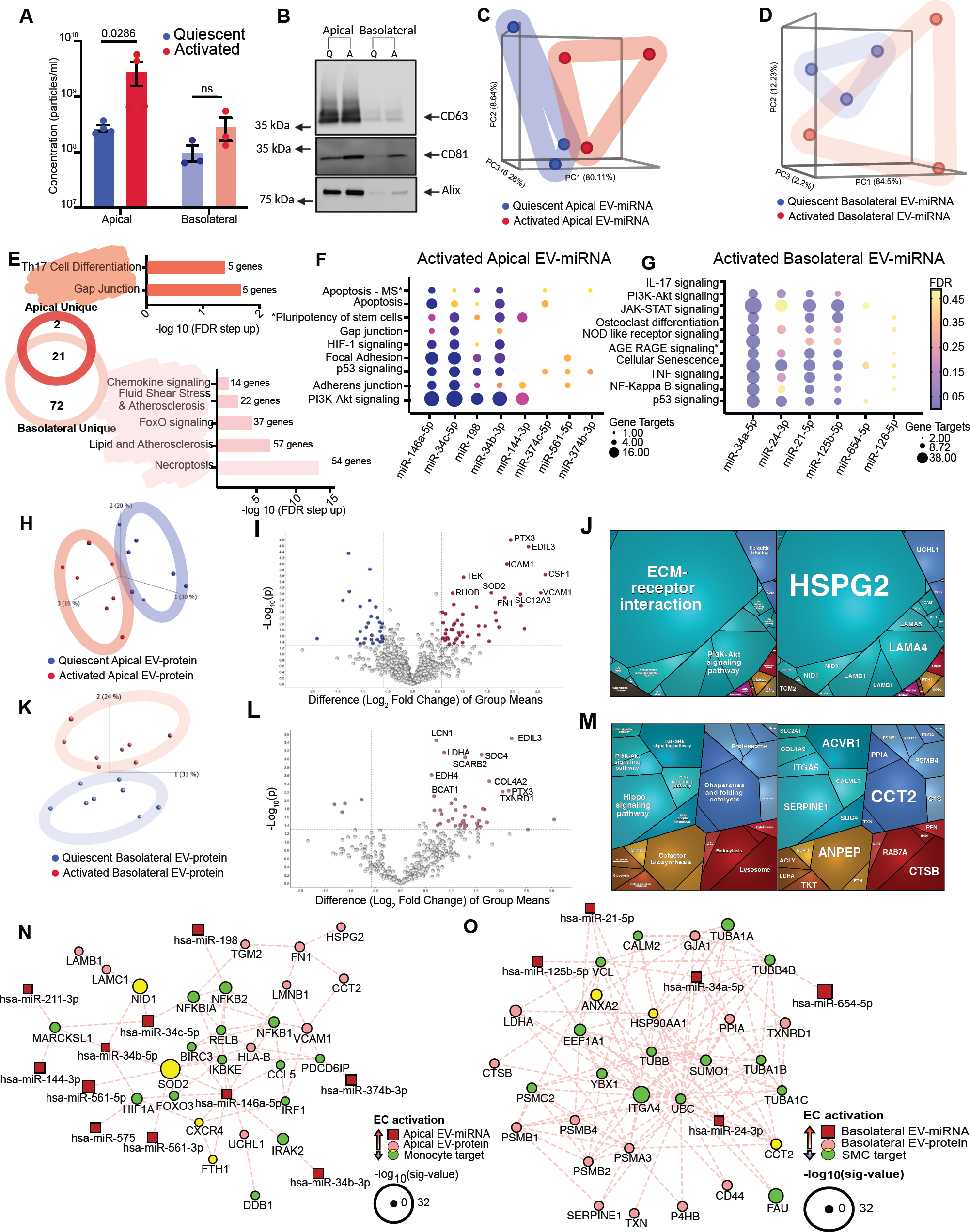
Activated endothelial cells modulate sEV miRNA and protein cargo in a compartment-specific manner with the capacity to uniquely affect circulating monocytes and resident vascular smooth muscle cells. **A-B,** Comparison of EC activation (red) versus quiescence (blue) on polarized sEV concentration in apical and basolateral compartments as determined by NTA **(A)** and western blot **(B)** (*n=*3-4). **B**, EV markers CD63, CD81, and Alix are denoted on the right, with molecular weights on the left. **C-D**, Unfiltered principal component analysis of apical (c) and basolateral EV-miRNA cargo **(D)** in activated versus quiescent states. E, KEGG pathway analysis (FDR<0.05) of top apical (n=10) and basolaterally (n=6) enriched miRNA highlight unique and shared pathways modulated by differentially expressed EV-miRNA in activated conditions (VENN diagram, left). Unique KEGG pathways enriched by activation in apical (dark red, top graph) and basolateral (light red, bottom graph) EC-EVs shown on right. Bar graph scaled by -log10(FDR) and labelled with number of genes involved in pathway. **F-G**, Activated versus quiescent EV-miRNA analyzed in apical **(F)** and basolateral **(G)** compartments. Top 10 EV-miRNAs (by FDR) were inputted for KEGG orthology pathway analysis with the addition of miRNA-146a-5p in apical conditions. KEGG pathways (FDR < 0.05) of interest showing individual miRNA associations. Data points are sized by GeneRatio (genes altered in pathway/total number of unique genes in analysis) and colour-scaled by FDR. **H-J**, EV-proteins enriched in the apical compartment by activated ECs. H, Unfiltered PCA analysis of apical EV-protein profiles in activated versus quiescent states (*n*=7). I, Volcano plot of differentially enriched EV proteins from the apical compartment in activation (red) and quiescence (blue) (p< 0.05, Fold Change |1.5|). Top ten differentially enriched proteins in the activated conditions are labelled. J, All differentially enriched proteins (p< 0.05, Fold Change |1.5|) from apical activated conditions were inputted to generate proteomaps (v2.0, Homo Sapiens), weighted by protein mass abundance. KEGG orthology terms (left) and respective proteins (right) contributing to the pathways are illustrated. K-M, EV-proteins enriched in the basolateral compartment by activated ECs. **K**, Unfiltered PCA analysis of basolateral EV-protein profiles in activated versus quiescent states. L, Volcano plot of differentially enriched EV proteins from the basolateral compartment as in **(I). M**, Proteomap (v2.0, Homo Sapiens) of differentially enriched basolateral EV proteins in activated conditions were inputted as in **(J). N**, Interactome integrating apical activated EC-EV secretome (top 15 miRNAs and all EV-proteins) with differentially expressed monocyte transcripts based on degree of interactions. O, Interactome integrating basolateral activated EC-EV secretome (top 15 miRNAs and all EV-proteins) with differentially expressed SMC transcripts based on degree of interactions. EV-miRNAs shown in red, EV-proteins in pink, and targets in green. Node size denotes significant value. *Signaling pathways regulating pluripotency of stem cells, AGE-RAGE signaling in diabetic complications.

Proteomics demonstrated that sEV cargo clustered by polarity (broad circles) with distinct proteomes in the apical versus basolateral compartment under quiescent conditions (Figure 5C), which was preserved with EC activation (Figure 5C and Online Figure VII D). While several known EV proteins were detected in all sEVs, regardless of compartment, we identified increased abundance in EV tetraspanins, EV-sorting and release proteins and intra-EV markers in apical sEVs, consistent with our observation that more sEVs are released apically (Online Figure VII C). There were 185 and 134 differentially enriched proteins in quiescent apical and basolateral EC-sEVs, respectively (Figure 5D, left panel). Endothelial sEV protein cargo from apical versus basolateral compartments reflected distinct pathways (Figure 5D, right panel). Apically derived sEV-proteins were associated with several metabolic pathways including glycolysis, protein metabolism, RNA metabolism, and maintenance of cell cytoskeleton, while basolateral sEV-proteins were predominantly associated with extracellular matrix interactions, cholesterol metabolism and transport, and protein degradation. There were several differentially enriched EV-trafficking proteins in both compartments (e.g., Apical: RAB7A, CAV1, EHD2,4; Basolateral: KRT2, LMNB1, EEA1, ANXA11), implicating them as potential mediators of selective and directional EV release (Online Figure VII C). Similar to the sEV-miRNA transcriptome, polarized release of sEV-proteins was maintained in activated states (Online Figure VII F).

Together, these data delineated the sEV miRNA and protein cargo released from ECs towards luminal and abluminal compartments. Under conditions of EC activation, polarization was preserved, but subtle secretome shifts (investigated below) were noted with expected consequences on cells that would typically receive EC-sEVs in the luminal circulation or abluminal vessel wall. The potential discovery of a mechanism underpinning endothelial communication with cells in different biological compartments that could be affected by an atheroprone stimulus prompted us to formally assess how EC activation alters compartment-specific sEV release, to directly compare sEV cargo between quiescent and activated states, and to test putative effects on monocyte and SMC recipients by *in silico* analysis.

### Luminal and abluminal cell reprogramming by polarized EC-EV cargo

Given that activated ECs have increased sEV release (Figure 1), we wanted to delineate whether this was a polarized phenomenon. Maintenance of a physiological barrier after IL-1β activation was similarly confirmed by VE-cadherin expression/localization and 30 nm gold nanoparticle challenge (Online Figure VI A, bottom panel; Online Figure VI B). While activated EC-sEVs from the apical or basolateral compartments appeared morphologically similar compared to quiescent EC-sEVs and were similar in size (Online Figure VIII A), IL-1β treatment significantly increased the concentration of EVs released into the apical, but not the basolateral, compartment (Figure 6A-B).

To determine the extent to which the polarized sEV-miRNA cargo profile changes when ECs are activated, we made direct comparisons between quiescence and IL-1β stimulation. sEV-miRNA clustered distinctly, according to activation status (Figure 6C-D, Online Figure VIII B-C). Under activated conditions, there were 23 and 93 pathways predicted by apical versus basolateral sEV-miRNAs, respectively. Many of these pathways were altered in basolaterally released EV-miRNA from activated ECs, emphasizing a previously undefined role for basolateral EC-sEVs participating in cell-cell communication within the vessel wall interstitium (Figure 6E). To that end, we delineated several key pro-atherogenic pathways that were uniquely predicted by basolaterally secreted sEV-miRNA from activated ECs, including chemokine signaling, response to shear stress, and lipid and atherosclerosis pathways. This implies a capacity for modulating focal vascular biology, as might be found in the atherosclerotic plaque microenvironment. Examining the miRNA specifically, well-known miRNA mediators of inflammation, miRNA-146a,^30^ miRNA-34c,^31^ miRNA-144,^28, 29^ and miRNA-374b^32^ were increased in apical EC-sEVs upon activation (Figure 6F). These sEV-miRNA were involved in several pathways previously predicted by the effects of plate-derived EC-sEVs on primary human monocytes (Figure 3C) such as apoptosis, cell proliferation and differentiation, and adhesion (Figure 6F). Levels of miRNA regulators of inflammation among the basolateral EVs such as miRNA-125b, miRNA-34a, miRNA-21, miRNA-24, and miRNA-126-5p were increased (Figure 6G). These sEV-miRNA likewise contributed to the pathways predicted by activated plate-derived EC-sEVs on SMCs (Figure 3H) including inflammatory signalling, cellular senescence, and cell proliferation and differentiation (Figure 6G).

We also detected distinct compartment-specific changes in the sEV-proteome with endothelial activation (Figure 6H-M, Online Figure VIII D). sEV-proteins clustered distinctly, according to activation status (Figure 6H,K). There were 47 sEV-proteins enriched in the apical compartment (Figure 6I) known to regulate pathways previously seen to be altered in monocytes treated with EC-sEVs (Figure 3), including proliferation (LAMA4/5, LAMB1, LAMC1, CSF1), inflammation (ICAM1, VCAM1, CSF), and cell adhesion and migration (LMNB1, HSPG2, NID1, NID2, CXCR4) (Figure 6J). While several of these pathways aligned with apically secreted sEV-miRNA (such as NF-kB and PI3K-Akt), response to oxidative stress via SOD2 secretion emerged as an sEV-protein specific function. As seen with the sEV-miRNA, the basolaterally secreted sEV-proteome from activated ECs predicted a diverse set of cellular functions further hinting at their important role in governing disease pathogenesis within the vessel wall (Figure 6L-M). There were 40 sEV-proteins enriched in the basolateral compartment from activated ECs (Figure 6L), many participating in pathways found predicted in EC-sEV-treated SMCs (Figure 3) including protein biosynthesis (BCAT, CCT2, PPIA, TXN, CCT2), inflammatory signalling (SLC2A1, CALML5, ACVR1), and PI3K-Akt signalling (COL4A2, ITGA5) (Figure 6M). Finally, our compartment-specific sEV interactomes integrating differentially expressed sEV-miRNA and sEV-protein from activated states further corroborated their role in inflammation, cell cycle, proliferation, and transcriptional and translational regulation (Online Figure VIII E-F). Though there was overlap between predicted targets of apical and basolateral EC-sEVs, endothelial activation altered sEV cargo that also led to distinct targets in each compartment with the basolateral interactomes emerging as denser networks.

Lastly, to discern luminal and abluminal endothelial sEV communication with relevant cell types in the context of an atheroprone stimulus, we generated *in-silico* interactomes by layering apical and basolateral EC-sEV cargo with the transcriptomic responses of sEV-treated monocytes and SMCs respectively (Figure 3), from activated conditions (Figure 6N-O and Online Figure VIII G). Apical sEV-miRNA and proteins enriched by endothelial activation (miR-146a, miR-198, miR-34b, miR-575, SOD2) altered monocyte mRNA transcripts involved in apoptosis (FOXO3, DDB1, PDCD61P, BIRC3), inflammatory signalling (NF-kB associated proteins, IRF1, IRAK2), adhesion and migration (MARCKSL1, NID1, CCL5), proliferation and differentiation (CCL5, CXCR4), and oxidative stress (HIF1A, SOD2) (Figure 6N). Basolateral sEV-cargo altered SMC mRNA transcripts known to function in activation-related pathways including protein biosynthesis (S and L ribosomal proteins, SUMO1, UBC, CCT2, FAU, PSMC2, EEF1A1), cell proliferation and differentiation (ITGA4, VCL, CALM2), cell senescence (HSP90AA1), and phagosomes (TUBB) (Figure 6O). Together, these data delineated polarized changes to apical and basolateral sEV-cargo upon EC activation with IL-1β and revealed that the unique shift in basolateral EC-sEV cargo is robust and affects athero-relevant pathways. Critically, our data suggest distinct roles for compartment-specific EC-sEV cargo on recipient cells found luminally in the circulation (e.g., monocytes) and abluminally within the vessel wall interstitium (e.g., SMCs) (see graphical abstract).

## Discussion

Here we demonstrate that ECs can utilize their ability to secrete sEVs directionally to communicate with luminal and abluminal cells in quiescent and activated states. This has important implications for cardiovascular diseases given the change in sEV cargo upon EC activation with IL-1β, a known mediator of inflammation in atherosclerosis and other cardiovascular pathologies. Primary human ECs secrete more CD63-positive sEVs upon activation and notably, this increase in sEVs is seen apically suggesting that circulating endothelial sEVs have the potential to serve as liquid biopsies of localized vascular disease (e.g., atherosclerotic plaques) where the endothelium is activated. Additionally, although the abundance of sEVs secreted basolaterally is not altered upon activation, we identify distinct changes in their cargo, with implications for cell-cell communication with cells in the vessel wall. ECs appear to load pro-inflammatory and atherogenic miRNAs and proteins into sEVs upon activation. Functionally, endothelial sEVs communicate with primary human monocytes and SMCs leading to changes in hundreds of protein coding transcripts, with unique responses depending on whether the endothelium is quiescent or activated. Our discovery that the endothelium is capable of directional sEV release provides a mechanism for focused communication with cells in discrete compartments. To that end, we found that ECs load starkly different sEV-cargo (miRNA and protein) for release apically versus basolaterally. Both apical and basolateral endothelial sEV content is altered upon activation with IL-1β, while *in silico* analysis underscored the ability for apical and basolateral messaging to alter transcripts in luminally and abluminally residing cells, respectively. This pronounced shift towards atherosclerosis pathways in basolateral sEV cargo after endothelial activation identifies a potential strategy for focal endothelial-based therapies in atherosclerotic disease.

Extracellular vesicles are known to be sentinels of disease states, and traffic biological entities between cells. ^33^ The quantity of EVs increase and their cargo is altered in several cardiovascular conditions.^34–44^ ECs are exquisitely sensitive to their surroundings and increase release of EVs in the presence of inflammatory mediators,^45–49^ and crucially, alter EV-contents (miRNA and protein) upon activation.^50–52^ We found that primary human ECs have increased CD63-positive sEV release when treated with IL-1β, a pro-inflammatory cytokine with known roles in mediating atherogenesis. Activated ECs secrete sEVs with miRNA and protein cargo that regulate key pathways involved in the pathogenesis of atherosclerosis including cell matrix interactions, cell-cell communication, cell death, protein synthesis, and inflammatory signalling.^53–56^ Interestingly, sEV-miRNA and protein cargo mediate varied functions in quiescent states, but the activated endothelial sEVs had denser networks and converged to modulate five key pathways in activated states: NF-kB signaling, lipid and atherosclerosis, necroptosis, cytokine-cytokine receptor interaction, and cell cycle. Biologically, monocytes treated with activated endothelial sEVs modulated pathways related to apoptosis, adhesion, migration, and proliferation, while SMCs demonstrated altered ribosomal and cellular senescence pathways. Together, sEV miRNA and protein cargo worked collectively to alter transcripts in recipient cells, stressing the importance of studying sEV functions in a holistic manner. The diversity in effects with the same effector is likely due, at least in part, to differences in the transcriptome of monocytes and SMCs, which alters the repertoire of miRNA targets available and cell signaling receptors that can respond to EV proteins. Although these data strengthened prior reports that ECs can communicate with surrounding cells via EVs^11, 12, 57^, it remained unexplored whether the endothelium might strategize polarized communication to cells in separate extracellular compartments.

Challenges exist for studying EV cargo selection.^58, 59^ However, the sheer number of ECs living at the interface of the circulation and blood vessel wall and potential for polarized sEV communication would inform several biologic processes. As a basic tenet, directional EV release necessitates a polarized structure capable of maintaining discrete compartments. In embryology, endothelial apical-basal polarity is a crucial component of angiogenesis with negatively charged glycoproteins concentrated at the apical surface to facilitate cord hollowing.^60, 61^ Proteins are segregated between the apical and basolateral surfaces in adult endothelial cells,^62–69^ with several EC-secreted proteins enriched luminally^70^ or abluminally^70–73^. While polarized EV release has been inferred from proteomics data,^70^ it has never been directly demonstrated, nor has the functional relevance of directional endothelial EVs been explored. We employed multiple approaches to demonstrate polarized secretion of sEVs by HAECs and confirmed our findings in another primary endothelium (HUVECs). In addition to imaging secreted sEVs from media (NTA and TEM), we imaged MVBs poised for release at both apical and basolateral surfaces (cryo-EM) and visualized basolateral release using TIRF microscopy. sEVs were quantified, profiled by miRNA and protein, and sEV interactomes generated. In quiescence, ECs released more CD63-positive sEVs apically and contained cargo involved in apoptosis, p53 signalling, metabolic pathways, and maintenance of cytoskeleton, while basolaterally secreted EC-EVs modulated pathways related to cell senescence, lipid and atherosclerosis, and cholesterol metabolism. The observation that ECs release distinct sEV miRNA and protein cargo from apical and basolateral surfaces shifts the current EC-EV communication paradigm, forcing a renewed consideration of cell-cell communication at this boundary region.

Given that ECs are frequently activated in cardiovascular disease, we chose to consider our findings in the context of atherosclerosis - the underlying cause of 19 million deaths globally per year.^74^ Upon activation with IL-1β, there were increased sEVs released apically and there were compartment-specific changes in the sEV-secretome with *in silico* evidence for compartment specific communication with luminal (monocytes) and abluminal cells (SMCs). Apically, the large pool of endothelial sEVs could represent systemic drivers of health and disease or could be useful as liquid biopsies that reflect vulnerable atherosclerotic plaques. Precedence for this exists in the cancer literature, where tumor derived EVs can drive metastatic disease^75^ and serve diagnostic roles.^75–79^ More striking however, was our novel finding that endothelial cells release sEVs basolaterally, and that activation leads to profound changes in the cargo released from the basolateral surface. This has implications for designing therapies that target atherosclerotic plaque biology in a focal manner. Emerging targets such as efferocytosis (a process for clearing dead cells that is defective in advanced plaques)^80–87^ or plaque stabilization through strengthening the SMC fibrous cap formation^88^ would be ideal. Alongside the cholesterol metabolism pathways enriched in basolateral sEVs, it is notable that efferocytosis-related proteins were seen basolaterally (LRP1, MFG-E8) but not apically (Online Figure VII F, right panel). If we can determine how to harness basolateral sEV release from ECs, it might be possible to deliver local plaque therapies targeting efferocytosis or cap-stabilization. To do so, future studies will need to target the activated endothelium, determine exactly how the endothelium designates specific cargo for loading, and to distinguish the intracellular trafficking pathways utilized for apical versus basolateral release.

Utilizing their capacity for polarized sEV release, ECs can participate in systemic and local cell communication. Capitalizing on this biology to modify endothelial-governed functions provides a powerful approach for detecting and/or modulating cardiovascular disease states such as atherosclerosis. The findings in this study provide early insights to support this exciting possibility. Together, these data provide a fresh perspective on endothelial sEV-based communication and the biological relevance of these messaging strategies in diseases such as atherosclerosis.

## Acknowledgements

The authors thank Dr. Alissa Weaver for kindly providing the pHluorin-CD63 plasmid (TIRF microscopy experiments) and Ms. Shiori Kuraoka for assistance with mass spectrometry. For invaluable discussions, we thank the following: Drs. Paul Fraser (TEM imaging interpretation), Shrey Sidwani (gold nanoparticle experiments), Dakota Gustafson (EV analysis), and Dr. Myron Cybulsky (critical discussion). This work was supported by Canadian Institutes of Health Research (CIHR) Project Grants PJT178006 (K. Howe) and PJT148487 (J. Fish), NIH grants (R01 HL136431, R01 HL141917, R01 HL147095; E. Aikawa), as well as a Tier II Canada Research Chair from CIHR (J. Fish). Additional support for this work was provided by the Heart and Stroke Foundation of Canada (New Investigator Award, K. Howe), Vascular Cures (Wylie Scholar Award, K. Howe), Blair Early Career Professorship in Vascular Surgery (K. Howe), Peter Munk Cardiac Centre (K. Howe), and University Health Network (K. Howe) and Vanier Canada Graduate Scholarship (S. Raju). Infrastructure funding was obtained from the John R. Evans Leaders Fund (J. Fish). T.W.W Ho is supported by a CIHR Canada Graduate Scholarship - Master’s Award. W.L.L. is supported by a Discovery Grant from the Natural Sciences and Engineering Research Council (NSERC) of Canada (RGPIN 2020-04299) and a Canada Research Chair in Mechanisms of Endothelial Permeability. Graphical abstract created with BioRender.com.

Conceptualization and data interpretation: SR, JEF and KLH. Writing original draft: SR and KLH. EV isolation, validation, and preparation for cargo interrogation: SR, KP, CC. EV transcriptomics: SR, KP. EV proteomics: SR, MB, CLC, TP, SS, EA. TIRF microscopy: TWWH, WL. CD63 plasmid transfection: SR, KP, RW. Electron microscopy: NLG, LF. Network analysis and EV interactomes: SR, SB, KLH. Manuscript editing: SR, SB, MB, CC, RW, SS, WL, EA, JEF, KLH. All authors read and approved the final manuscript. Funding acquisition: JEF and KLH. Supervision of the study: KLH.

---

**Online Figure I.**
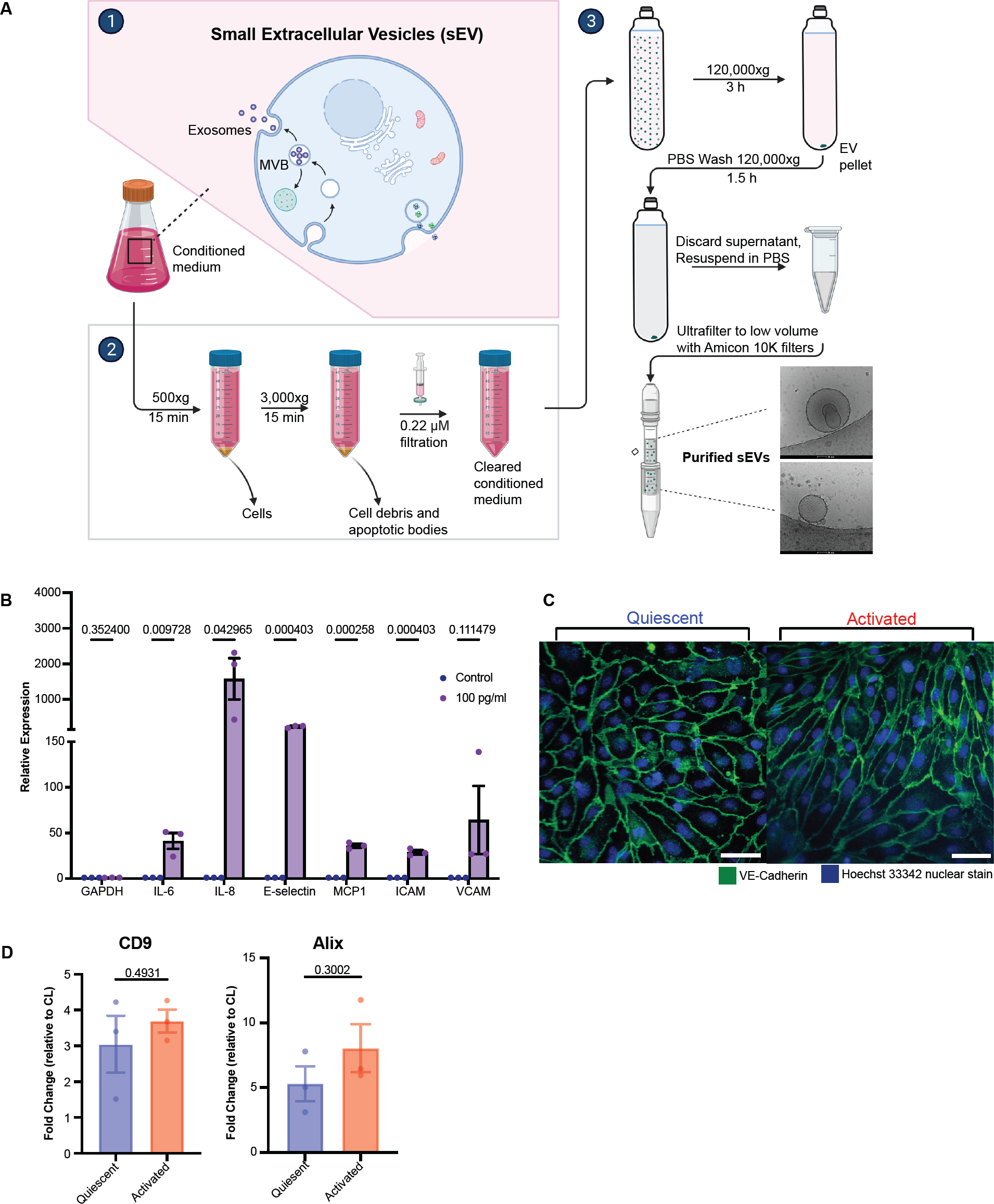
General features of sEV isolation in human aortic endothelial cells in quiescence and after confirmed activation with IL-1β. **A,** Schematic for sEV enrichment. Endothelial cells were grown to confluence and maintained in EV-free media for 24 h prior to supernatant collection. Conditioned media was centrifuged at 500xg and 3,000xg for removal of cell debris and apoptotic bodies, followed by filtration with 0.22 μM to generate cleared conditioned media. EVs are enriched via ultracentrifugation at 120,000×g for 3 h, followed by a PBS wash, and ultrafiltration using a Amicon 10 kDa filter. EV enrichment was confirmed as per the MISEV2018 guidelines. Created with BioRender.com. **B,** RT-qPCR of inflammatory cytokines and adhesion molecules in cultured HAECs post treatment with 100 pg/mL IL-1β, 24 h. mRNA abundance was normalized to GAPDH. **C,** HAECs were grown on coverslips, placed in EV-free media (left) +/- IL-1β (right;100 pg/mL, 24 h) and stained for the adherens junction, VE-Cadherin (*n*=3). **D,** Densitometry of EV lysate derived CD9 and Alix normalized to HAEC cell lysate control. Bar graphs show mean ± SEM. Statistical significance assessed by multiple unpaired t-test with adjustment for multiple testing with the Benjamini-Hochberg procedure.

**Online Figure II.**
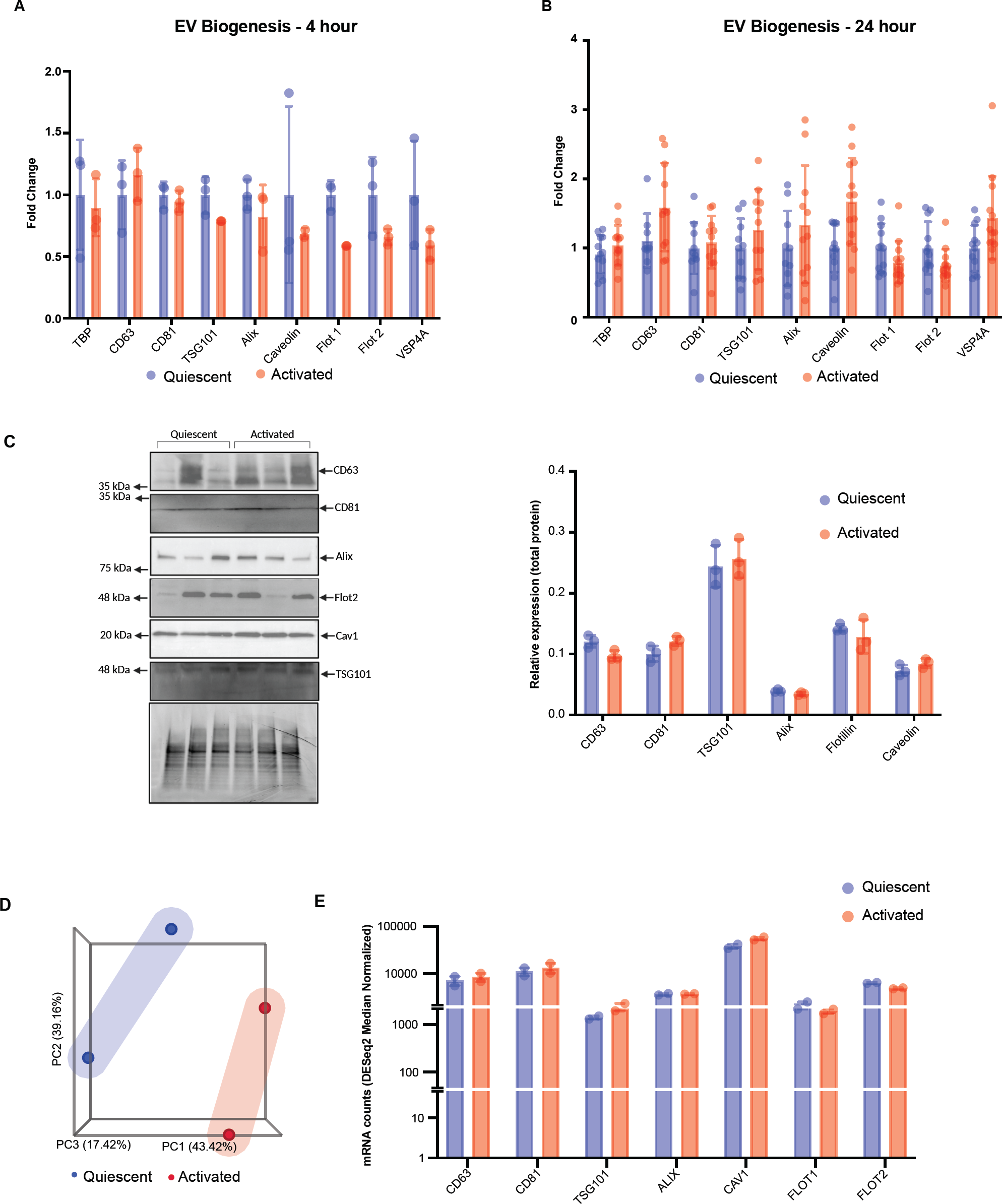
sEV biogenesis is unaffected by endothelial activation. **A-B,** RT-qPCR of genes known to function in EV sorting and release in cultured HAECs post treatment with 100 pg/mL IL-1β at 4 h **(A)** and 24 h **(B)**. mRNA abundance was normalized to the housekeeping gene, TBP. **C,** Western blot depicting expression of proteins involved in EV sorting and release in HAEC cell lysate (left). Densitometric analysis of EV markers, normalized to total protein (right). Arrows show position of correct protein band and molecular weights markers indicated on left. **D-E,** Publicly available HAEC RNA-seq data (GEO accession: GSE89970). HAECs were isolated from aorta of adult patients and activated with IL-1β (10 ng/mL, 4 h). **D,** PCA analysis. **E,** Median Ratio normalized mRNA counts of EV biogenesis proteins. Bar graphs show mean ± SEM. Statistical significance assessed by multiple unpaired t-test with adjustment for multiple testing with the Benjamini-Hochberg procedure.

**Online Figure III.**
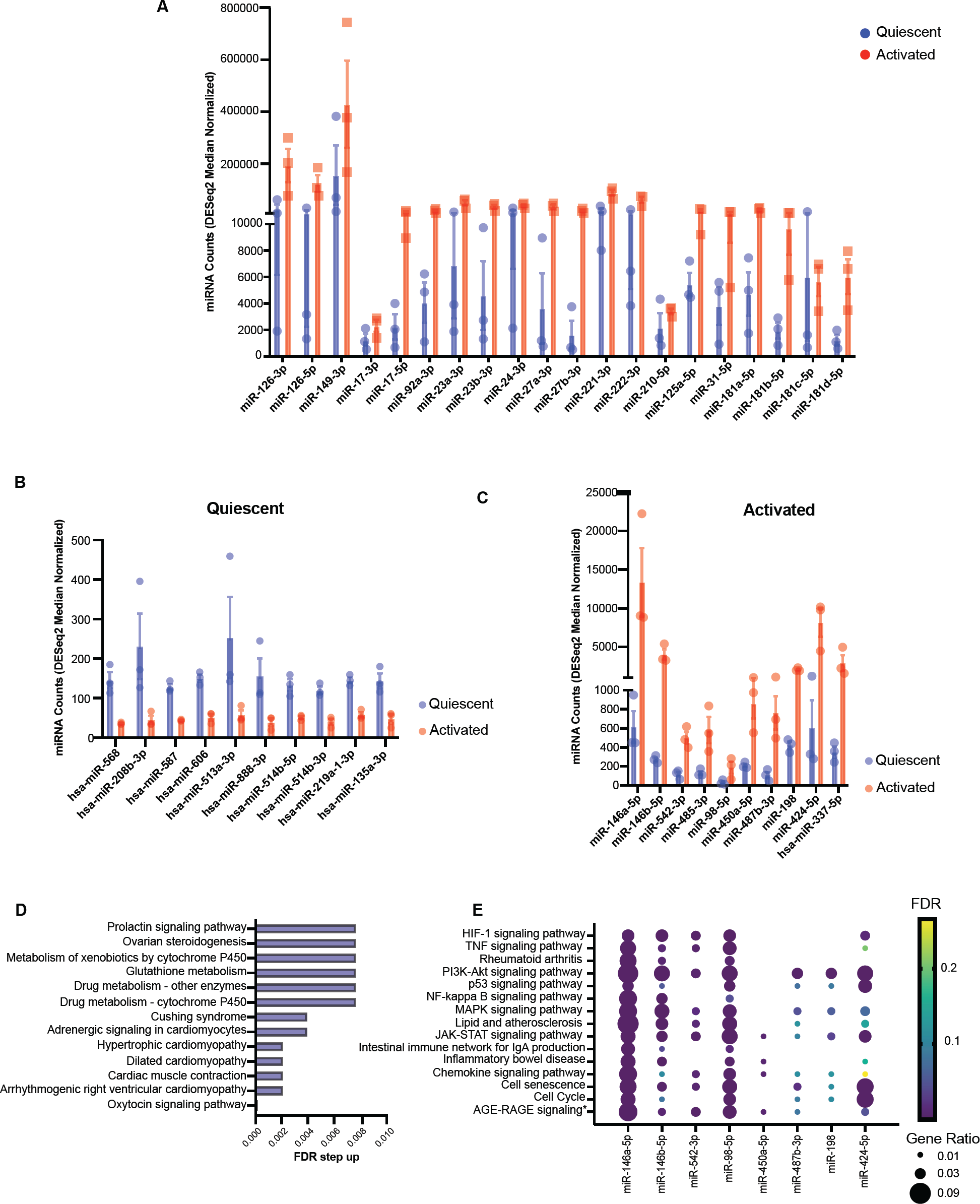
Additional analysis of sEV miRNA cargo in quiescent and activated endothelium. **A,** Median normalized miRNA counts of endothelial enriched miRNA in quiescent (blue) and activated (red) states. **B,** Median normalized miRNA counts of quiescent enriched EV-miRNA used in KEGG pathway analysis. **C,** Median normalized miRNA counts of activation enriched EV-miRNA used in KEGG pathway analysis. **D-E,** KEGG pathway analysis with top FDR-based pathways of EV-miRNA enriched in quiescent **(D)** and activated **(E)** states. Data points are sized by GeneRatio (genes altered in pathway/total number of unique genes in analysis) and colour-scaled by FDR.

**Online Figure IV.**
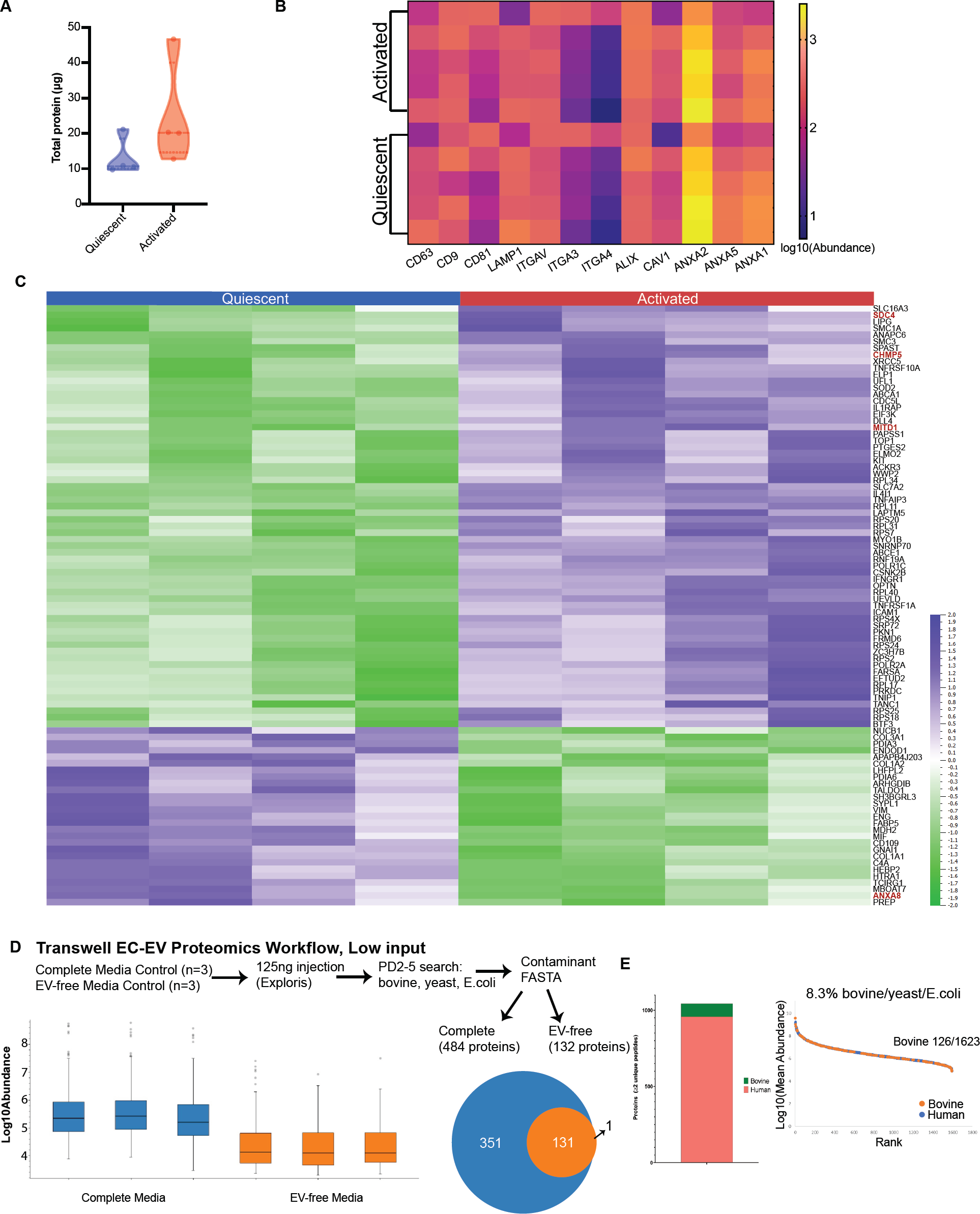
Workflow and quality control for sEV proteomics from quiescent and activated endothelial cells. **A,** Total protein quantification in HAEC EV lysates from quiescent and activated conditions. **B,** Abundances of common EV protein in HAEC EV-enriched samples in quiescent and activated conditions (CD63, CD9, CD81, LAMP1, ITGAV, ITGA3, ITGA4, ALIX, CAV1, ANXA2/5/1). **C,** Differentially expressed proteins in activated versus quiescent conditions (p≤ 0.05, Fold Change |1.5|). Proteins involved in EV trafficking and release are labeled in red. **D,** Assessment of media-based contamination for EC-EV transwell proteomics. Workflow shown on top. Normalized peptide abundances in complete media and EV-free media (left). VENN diagram depicting total number of proteins with two unique peptides in complete versus EV-free media (right). **E,** Quantification (left) and rank plot (right) of total proteins (with 2 unique peptides) derived from serum (labelled bovine) in our EV-samples.

**Online Figure V.**
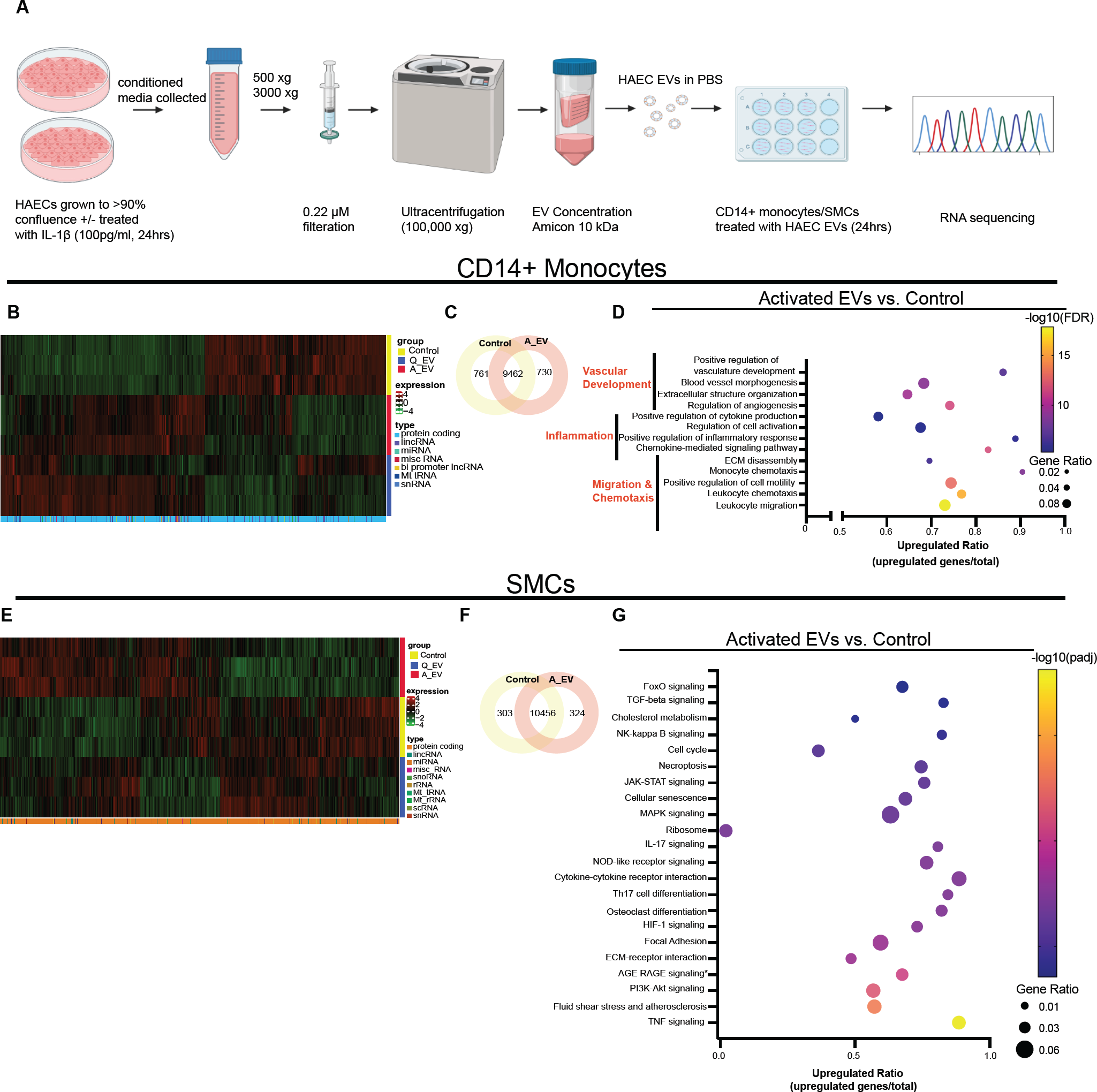
Effects of endothelial sEVs on recipient monocytes and smooth muscle cells. **A,** Schematic of experimental design. HAECs (+/- IL-1β treatment 100 pg/mL, 24 h), conditioned media collected, cell debris removed via centrifugation, filtered for sEVs, isolated by ultracentrifugation, and concentrated until resuspension and addition to primary human CD14+ monocytes (10^9–10^ sEVs added to 500,000 monocytes) or SMC (10^9–10^ sEVs added to 400,000 SMCs). After 24 h sEV exposure, monocyte cell lysates were collected, RNA isolated, purity confirmed by BioAnalyzer, and sent for RNA sequencing (400 ng, Novogene). **B,** Unfiltered heatmap analysis showing transcript abundance in treatment groups. Shading represents expression levels. Right legend identifies treatment group. Bottom legend identifies transcript type (protein coding vs. non-coding). **C,** VENN diagram depicting number of shared and unique monocyte RNA transcripts in comparisons of activated vs control groups. **D,** GO pathway analysis of the effects of activated EC-EVs versus media control on the monocyte RNA transcriptome (adjusted p-values ≤0.05 and |log2(FoldChange)| > 0). Data points are sized by GeneRatio (genes altered in pathway/total number of unique genes in analysis) and colour-scaled by FDR. Upregulated ratio was calculated by dividing the number of upregulated genes by the total number of genes known to function in each pathway. **E,** Unfiltered heatmap analysis showing transcript abundance in treatment groups as in **(B). F,** VENN diagram depicting number of shared and unique SMC RNA transcripts in comparisons of activated vs control groups. **G,** KEGG pathway analysis of the effects of activated EC-EVs versus media control on the SMC RNA transcriptome (adjusted p-values ≤0.05 and |log2(FoldChange)| > 0). Data visualization completed as in (**D).**

**Online Figure VI.**
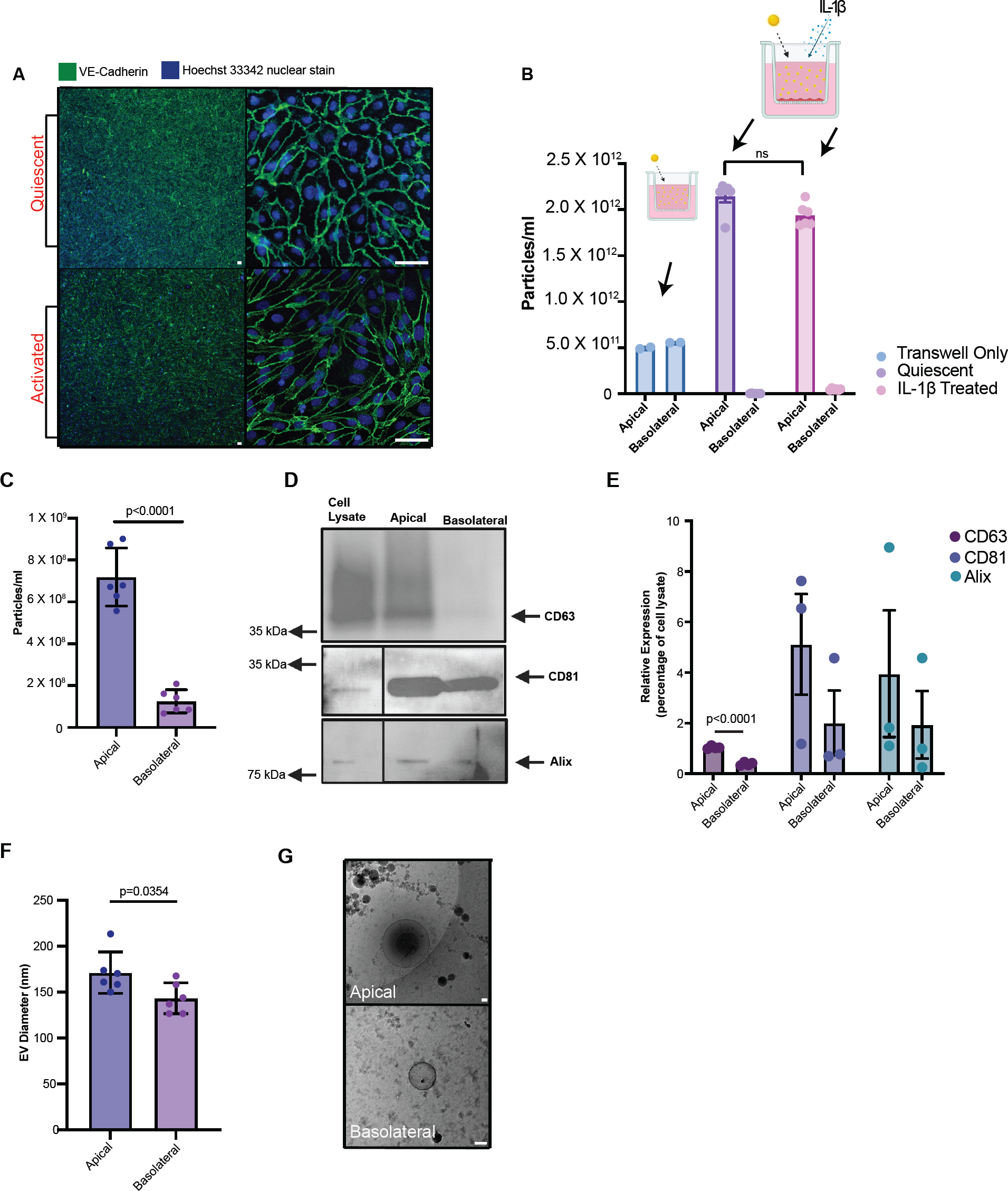
Validation of the model for polarized sEV release from endothelial monolayers. **A-B,** Endothelial cell physiologic barrier demonstration by VE-cadherin expression **(A)** and 30 nm gold nanoparticle challenge **(B). A,** HAECs were grown on transwell supports as described above, placed in EV-free media (top panel) +/- IL-1β (bottom panel;100 pg/mL, 24 h) and stained for the adherens junction, VE-Cadherin. **B,** Gold nanoparticle assay confirming the smallest EV-like nanoparticle (30 nm) does not cross the EC monolayer in quiescence or after activation with IL-1β at 100 pg/mL. **C-G,** HUVECs confirm polarized release of EVs to apical and basolateral compartments. **C,** Nanoparticle tracking analysis quantifying concentration of EC-EVs in apical and basolateral compartments. **D-E,** Western blot depicting protein expression of EV markers (positive (CD63, CD81, Alix), in cell lysate, apical EV and basolateral EV samples **(D)**. Arrows show position of correct protein band and molecular weights markers indicated on left. Densitometric analysis of EV markers **(E). F,** Nanoparticle tracking analysis quantifying the mean EV diameter in apical and basolateral compartments. **G,** Cryo-EM of representative images of apical (top) and basolateral (bottom) quiescent EC-EVs. Scale bar=50 nm.

**Online Figure VII.**
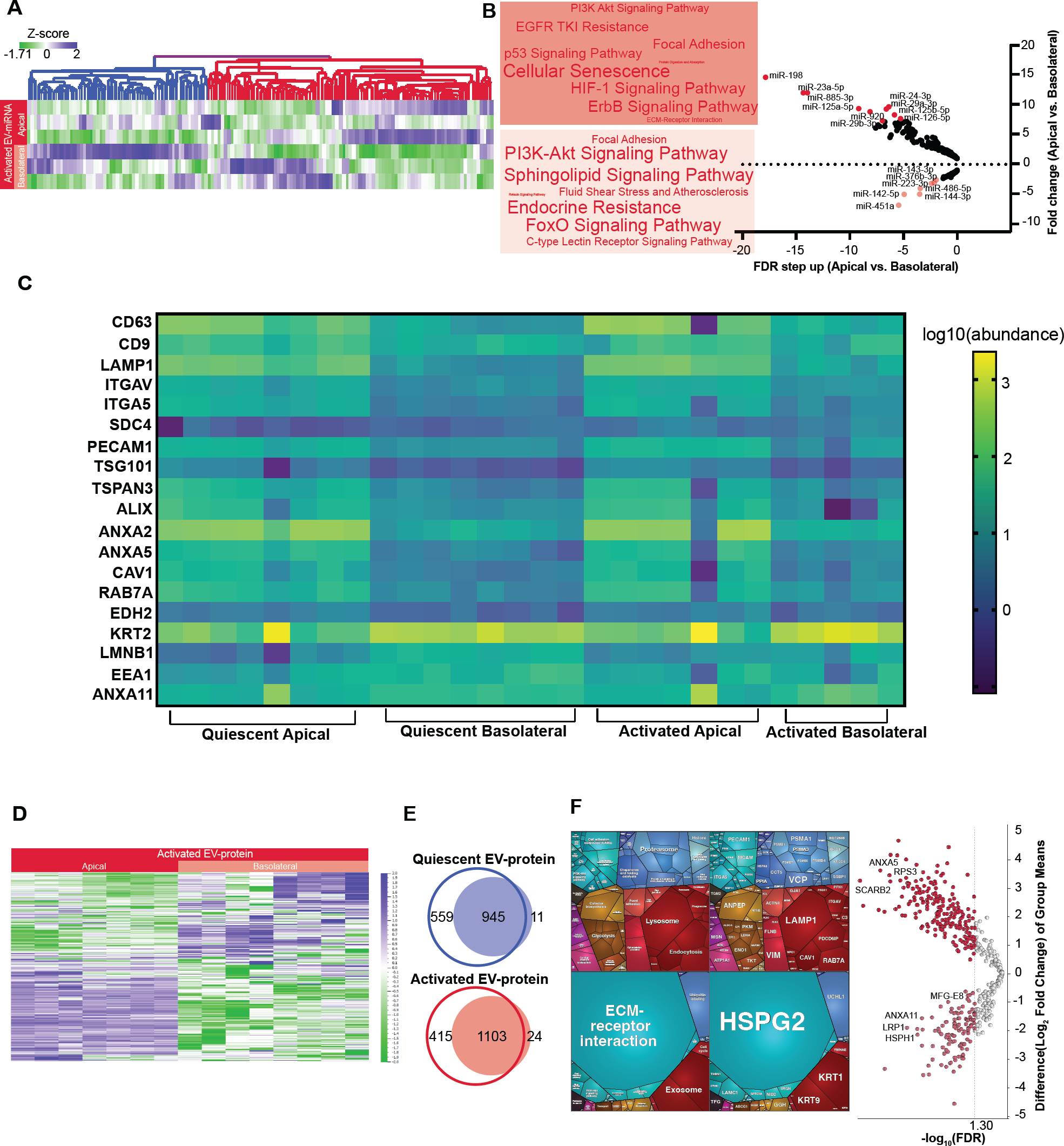
The phenomenon of polarized sEV release with distinct apical and basolateral miRNA and protein cargo is preserved after endothelial cell activation. **A-B,** Differential expression of EV-miRNA in apical versus basolateral compartments as depicted by unfiltered heatmap analysis (A) and KEGG pathway analysis **(B). B**, Volcano plot (right panel) of activated HAEC secreted EV-miRNA transcriptome enriched in apical (dark shading) versus basolateral (light shading) compartments. Top miRNA, by FDR step up, are labelled in each condition and used for downstream pathway analysis (FDR step up ≤ 0.05). KEGG pathway analysis of labelled miRNA in each condition (FDR ≤ 0.05), weighted by number of miRNAs participating in each pathway depicted by Word Cloud (left panel). C, Heatmap depicting EV protein markers (derived from EV proteomics) in apical and basolateral compartments for both quiescent and activated states. **D-F**, EV-proteomic analysis comparing apical versus basolateral EV-proteins. **D**, Unfiltered heatmap analysis depicting protein abundances of apical and basolateral EC-EVs from activated states (*n*=5-8). **E**, VENN diagrams depicting number of shared and unique proteins in comparisons of apical (open circle) versus basolateral (filled circle) in quiescent (top) and activated (bottom) states. **F**, Volcano plot (right panel) of activated HAEC secreted EV-proteome enriched in apical (dark shading) versus basolateral (light shading) compartments (FDR≤0.05). Left panel: All differentially enriched proteins (FDR≤0.05) in activated conditions were used to generate proteomaps (v2.0, Homo Sapiens), weighted by protein mass abundance. Apical and basolateral proteomaps are represented by top and bottom panels, respectively. KEGG orthology terms (left) and respective proteins (right) contributing to the pathways are illustrated.

**Online Figure VIII.**
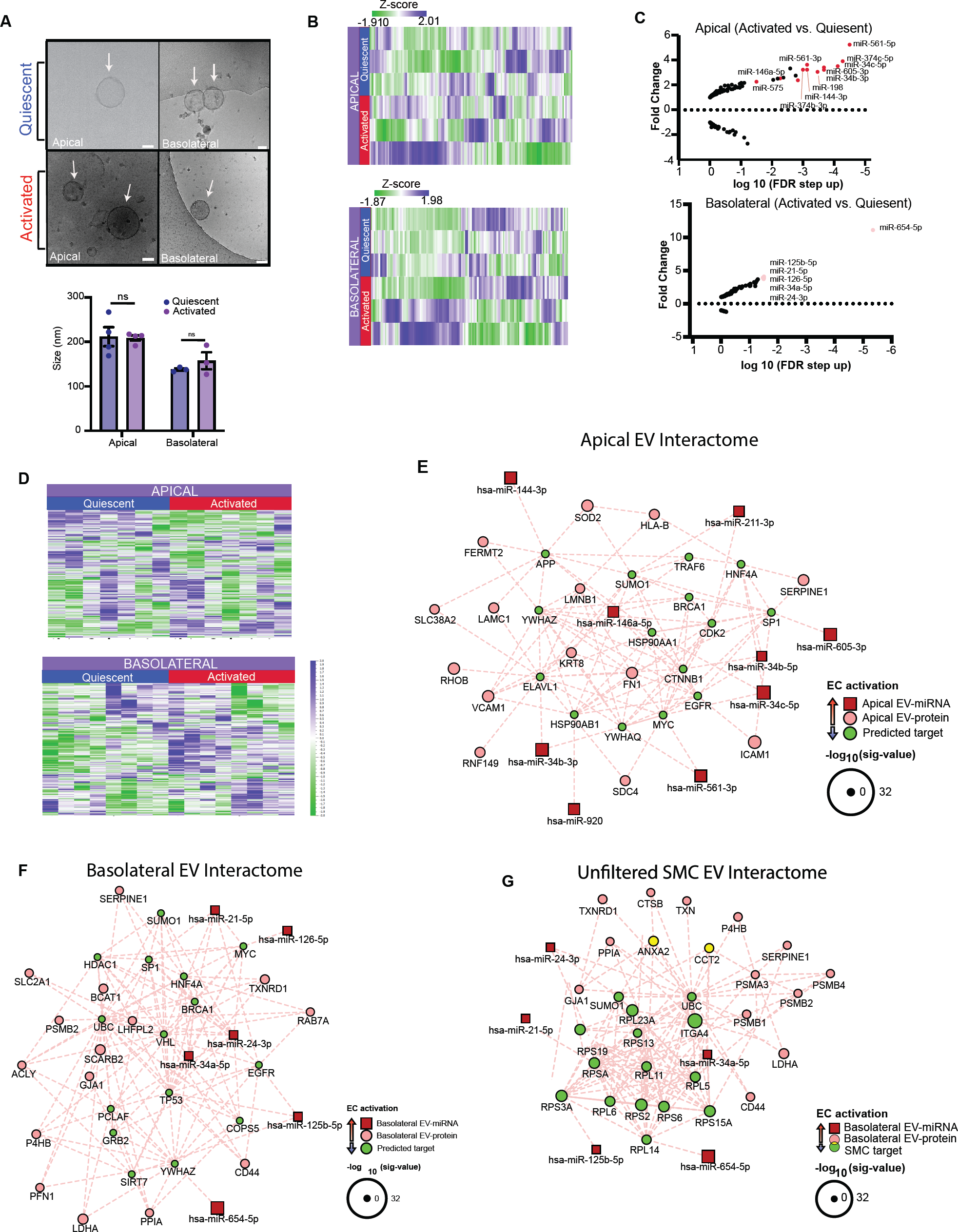
Comparison of quiescent and activated endothelial sEV cargo by biological compartment. **A,** Cryo-EM images of apical (left panel) and basolateral (right panel) sEVs released by quiescent (top) and activated (bottom) ECs. **B-C,** Comparison of activated versus quiescent EV-miRNA cargo in apical and basolateral sEVs. **B,** Unfiltered heatmaps of endothelial EV-miRNA transcriptome clusters by activation state in apical (top) and basolateral (bottom) sEVs. **C,** Volcano plots of activated versus quiescent HAEC secreted EV-miRNA transcriptome enriched in apical (top) and basolateral (bottom) compartments (FDR<0.05). **D,** Unfiltered heatmaps depicting EV-protein abundances in activated versus quiescent ECs by compartment (apical, top; basolateral, bottom). **E-F,** Interactomes of polarized EV release in activated conditions generated by capturing differentially expressed EV-miRNA (top 25 by FDR) and all EV-proteins, followed by network reduction to retain the top 15 of each group based on degree of interactions. **E,** Apical EV interactome with predicted targets. **F,** Basolateral EV interactome with predicted targets. **G,** Interactome integrating basolateral activated EC-EV secretome (top 15 miRNAs and all EV-proteins) with differentially expressed SMC transcripts based on degree of interactions and with inclusion of ribosomal targets. EV-miRNAs shown in red, EV-proteins in pink, and targets in green. Node size denotes significant value.

## NOVELTY AND SIGNIFICANCE

### What is known?

- Endothelial cells (ECs) are activated in regions prone to forming atherosclerosis.
- ECs release extracellular vesicles (EVs) in quiescent and activated states, implicating EC-EVs as a potential vector of cell-cell communication.
- Upon uptake by recipient cells, EVs can modulate biological processes via their miRNA and protein contents.

### What new information does this article contribute?

- ECs are dynamic and respond to pro-atherogenic stimuli by increasing release of a specific population of small EVs (sEVs) and altering their microRNA and protein cargo.
- sEVs released by ECs differentially reprogram key vascular cells such as primary human monocytes and smooth muscle cells (SMCs) towards an athero-prone signature.
- ECs release sEVs bidirectionally in quiescent and activated states, with distinct cargo capable of reprogramming recipient cells located in discrete vascular compartments.

The endothelium is a single layer of cells lining every blood vessel that lives at the interface between two dynamic environments. Activated ECs release more sEVs, which carry altered microRNA and protein cargo capable of driving atherosclerosis. EC-EVs communicate with monocytes and SMCs - cells predominantly contained in the circulation and vessel wall respectively - leading to changes in hundreds of protein coding transcripts, with unique responses depending on whether the endothelium is quiescent or activated. ECs are capable of directional communication through their ability to release EVs bidirectionally and by directing distinct cargo to apical (circulation) and basolateral (vessel wall) compartments. Both apical and basolateral endothelial sEV content is altered upon activation, and *in silico* analysis underscored the ability for apical and basolateral sEV messaging to alter transcripts in luminally and abluminally residing cells, respectively. Activated/inflamed ECs release more sEVs with atheroprone cargo apically; however, it is the basolaterally released sEVs that demonstrate the most profound shift in EV cargo towards athero-prone and inflammatory pathways implicating them as critical players in plaque biology. Together, these findings conceptually advance our understanding of EC cell-cell communication. Harnessing bidirectional endothelial sEV release represents a new frontier in diagnostics and therapeutics for cardiovascular disease.

